# A Notch signal required for a morphological novelty in *Drosophila* has antecedent functions in genital disc eversion

**DOI:** 10.1101/2025.05.09.653167

**Authors:** Donya N. Shodja, William J. Glassford, Gavin Rice, Sarah J. Smith, Mark Rebeiz

**Author notes:** Department of Biological Sciences, The George Washington University, Washington, DC 20052, USA. Department of Biochemistry and Molecular Biophysics, Columbia University, New York NY, 10027 USA. Department of Molecular Cellular and Developmental Biology, Yale University, New Haven CT, 06511, USA. Corresponding author: Mark Rebeiz **Email:**.

## Abstract

The origin of morphological novelties has long fascinated biologists. Signaling pathways play important roles in the formation of novelties, however, the history of how they become integrated into new developmental programs remains unclear. Here, we investigated the evolution of the posterior lobe, a novel structure in the male genitalia of *Drosophila melanogaster*. We demonstrated that a Notch signaling center is required for the formation of this novelty, and identified enhancers of the ligand *Delta*, which allowed us to track the evolutionary history of this signaling center. Surprisingly, we found that the posterior lobe signaling center emerged from a pre-existing role in genital disc eversion. We provide a likely mechanism by which Delta contributes to genital eversion through a network of apical extracellular matrix, which also became integrated into the posterior lobe program. This work demonstrates that novelties may be formed in the context of already complex developmental processes, by appending new roles to pre-existing signals.

## Introduction

The evolutionary origin of new morphological structures (“morphological novelties”) remains an enigmatic process that has captivated the imagination of biologists for centuries. Despite astounding anatomical diversity in the animal kingdom, genes that govern the formation of novelties are often conserved even between distantly related taxa (1–3), and recurrently used in multiple tissues over the course of an organism’s development (4,5). Specifically, a core set of signaling pathways are frequently re-used throughout development to regulate the formation of different tissues (6,7). Though elegant work on complex morphological novelties such as the turtle’s shell (8,9), butterfly eyespots (10,11), the bat’s wing (12–14), and birds’ feathers (15) have implicated a prominent role of signaling pathway re-deployment in their origination, we currently lack a molecular picture of how these roles first appeared and were shaped into complex genetic programs. In particular, we need detailed histories of how important developmental signals became expressed to pattern these structures and how downstream cellular responses first emerged.

To study the origination of a novel structure, two factors have hindered progress towards developing a satisfying molecular history of how novelties were assembled. First, many novel morphologies of interest have evolved over long evolutionary times, such that the ancestral context from which the novelty first emerged cannot be traced with any confidence (9,16). Indeed, some developmental evolutionary studies of novelty have focused on the development of single organisms without comparing them to outgroup species that lack the structure (9,10,17,18). This is an understandable approach for macroevolutionary changes which arose in the distant past, as conserved landmarks to reference homologous tissues are often absent. However, focusing on ancient novelties may overlook multiple layers of change that erased informative intermediate steps along their evolutionary trajectories. Thus, evaluating novelties across a wide range of timescales may offer multiple unique insights concerning their beginnings. A second barrier is that most systems lack the genetic and developmental tools necessary to assess gene regulatory changes and the function of genes in the relevant tissue (8,19,20). Both of these barriers have necessitated the study of more recently evolved morphological novelties that exist in genetically tractable model organisms (21–30). Doing so allows us to assess and pinpoint changes within the gene regulatory networks underlying these novelties.

Gene regulatory networks (GRNs) control development by integrating spatial and temporal information from transcription factors and signaling pathways to pattern the expression of target genes (31). In the context of morphological novelty, the study of enhancers uniquely provides access to study mutations that may have contributed to the evolution of these traits through comparative reporter assays (22,28). By evaluating the activities of orthologous regulatory regions from two or more species in a common genetic background, one can attribute activity differences to the tested regulatory elements. Furthermore, reporter assays can detect co-option and pleiotropy (22,32,33). When a program downstream of a signaling pathway is deployed to a new context, the responsible enhancer will be pleiotropic for both ancestral and novel tissues (22). Finally, enhancer elements are particularly useful for tracing cellular lineages through complex morphogenetic movements that may be difficult to disentangle.

The genitalia of the model organism *Drosophila (D.) melanogaster* offers a system in which a relatively recent morphological novelty can be examined in a genetically tractable system. Genital traits are noteworthy for their rapid evolution (34) and are often the distinguishing characteristic between closely related species (35). Notably, the posterior lobe is a morphological novelty in the male genitalia of *D. melanogaster* (35,36) (Fig. 1A-B). This cuticular outgrowth is unique to the *melanogaster* clade and is necessary for copulation (35,37,38). The posterior lobe projects from an ancestral genital tissue called the lateral plate (also known as the epandrial ventral lobe (39)) and forms by locally increasing the apical height of lateral plate cells to generate the lobe (40). This structure evolved approximately 35 million years ago (41), and the genitalia of lobed and non-lobed species are composed of similar structures otherwise. This permits us to perform comparative analyses of gene expression and gene regulation during development between *D. melanogaster* and outgroup species that lack a lobe, such as *D. ananassae* and *D. biarmipes*, which serve as a developmentally accessible proxy for the ancestral ground state (Fig. 1C, Fig. S1).

**Fig. 1.**
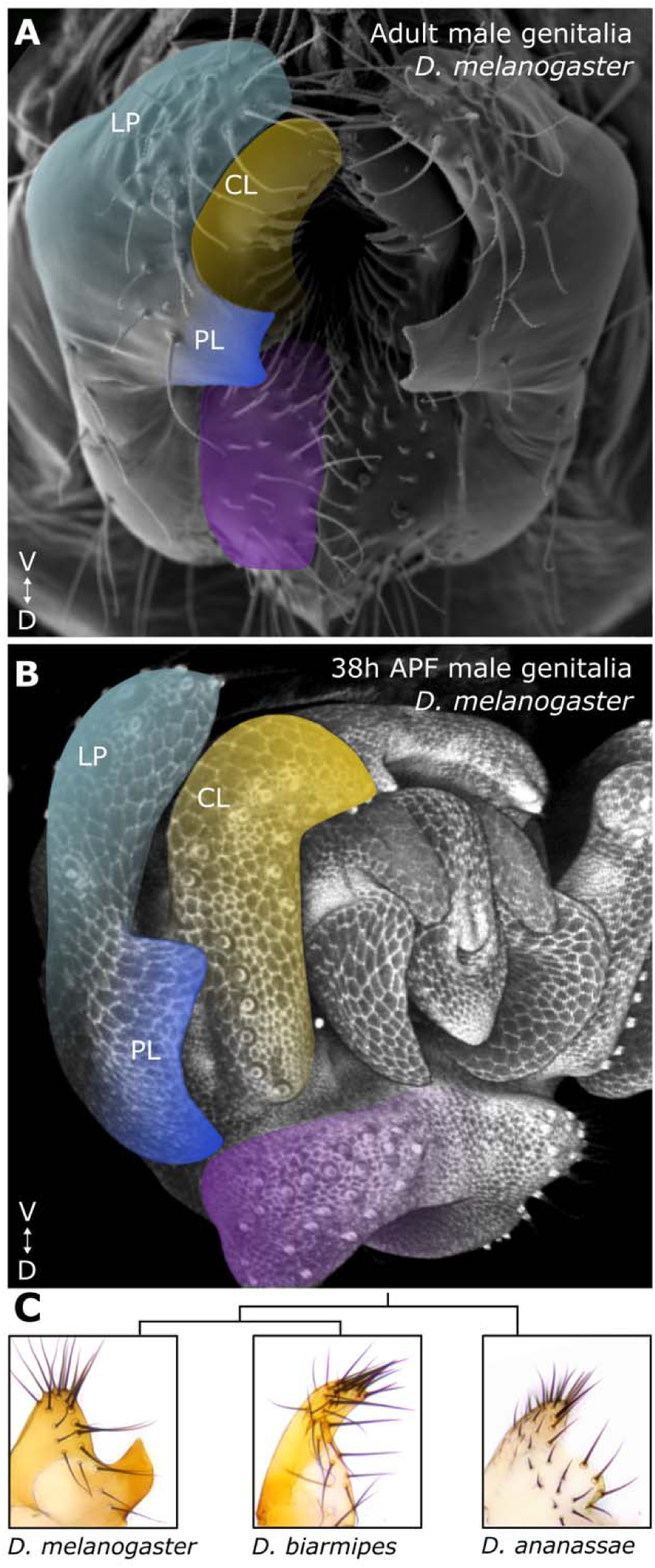
Male genitalia of *Drosophila melanogaster.* *D. melanogaster* adult **(A)** and developing pupal (38h APF) **(B)** male genitalia. The novel posterior lobe (royal blue) protrudes from an ancestral structure called the lateral plate (light blue), adjacent to another structure conserved between lobed and non-lobe species, the clasper (yellow). The analia is highlighted in purple. **(C)** Cladogram with brightfield images of dissected adult lateral plates of a lobed and two non-lobed species.

Here, we investigated how a signaling pathway became associated with the novel posterior lobe. We found that Notch signaling plays an important role for posterior lobe development. Specifically, the spatial expansion of the Notch ligand, Delta, in a zone adjacent to the posterior lobe is required for its development, and this expansion is unique to *D. melanogaster*. We dissected the regulatory elements involved in the deployment of Delta to the lobe-forming region and investigated its ancestral function in the development of genital structures. Surprisingly, our analysis revealed that the Delta/Notch signaling center becomes active days before the posterior lobe forms, serving a role in the development of conserved genital tissues. In particular, we’ve implicated an early-acting role for this signal in controlling genital disc eversion – a process by which the epithelium underlying these structures turns inside out. This work demonstrates that novelties may be formed in the context of ancestrally complex developmental programs, by adding new roles to pre-existing signals to connect a new program to well established ancestral ones.

## Results

### Species-specific expression of *Delta* is essential for development of the posterior lobe

In a screen of major signaling pathway ligands during posterior lobe development, we found that the Notch ligand, Delta, is expressed in multiple male genital structures, including a region adjacent to the developing posterior lobe at the base of the lateral plate and clasper (also known as the surstylus (39)) (Fig. 2A and Fig S2A-C). Importantly, this expression precedes posterior lobe development in the early pupal genitalia, as expected for a developmentally instructive signal (Fig. S2A). As the posterior lobe initiates its development at mid-pupal stages, the expression pattern of Delta spatially expands along the lateral plate and clasper boundary (Fig. S2B). Once the posterior lobe forms, Delta’s expression retracts dorsally towards the anal plate (Fig. S2C). The early expression of Delta preceding posterior lobe development and its expansion corresponding to the developmental timing of the posterior lobe made it a strong candidate regulator of the posterior lobe gene regulatory network. To determine whether this pattern is unique to lobe-bearing species, we next examined the expression of Delta in *Drosophila* species that do not form a posterior lobe (“non-lobed” species). Immunofluorescent staining of Delta using a polyclonal antibody that is cross-reactive in multiple species, as well as *in situ* hybridization of *Delta* mRNA, revealed that Delta is expressed in a small area at the base of the claspers and lateral plates in the non-lobed genitalia of *D. biarmipes* and *D. ananassae* (Fig. 2E-F, Fig S2D-F). This expression pattern is limited to a much smaller zone compared to *D. melanogaster* (Fig. 2D-F and Fig. S2D-F and Fig. S3G-I and J-L). These results suggested that the expansion of Delta is specific to the posterior lobe forming species, *D. melanogaster*, and correlates with the timing of posterior lobe development.

**Fig. 2.**
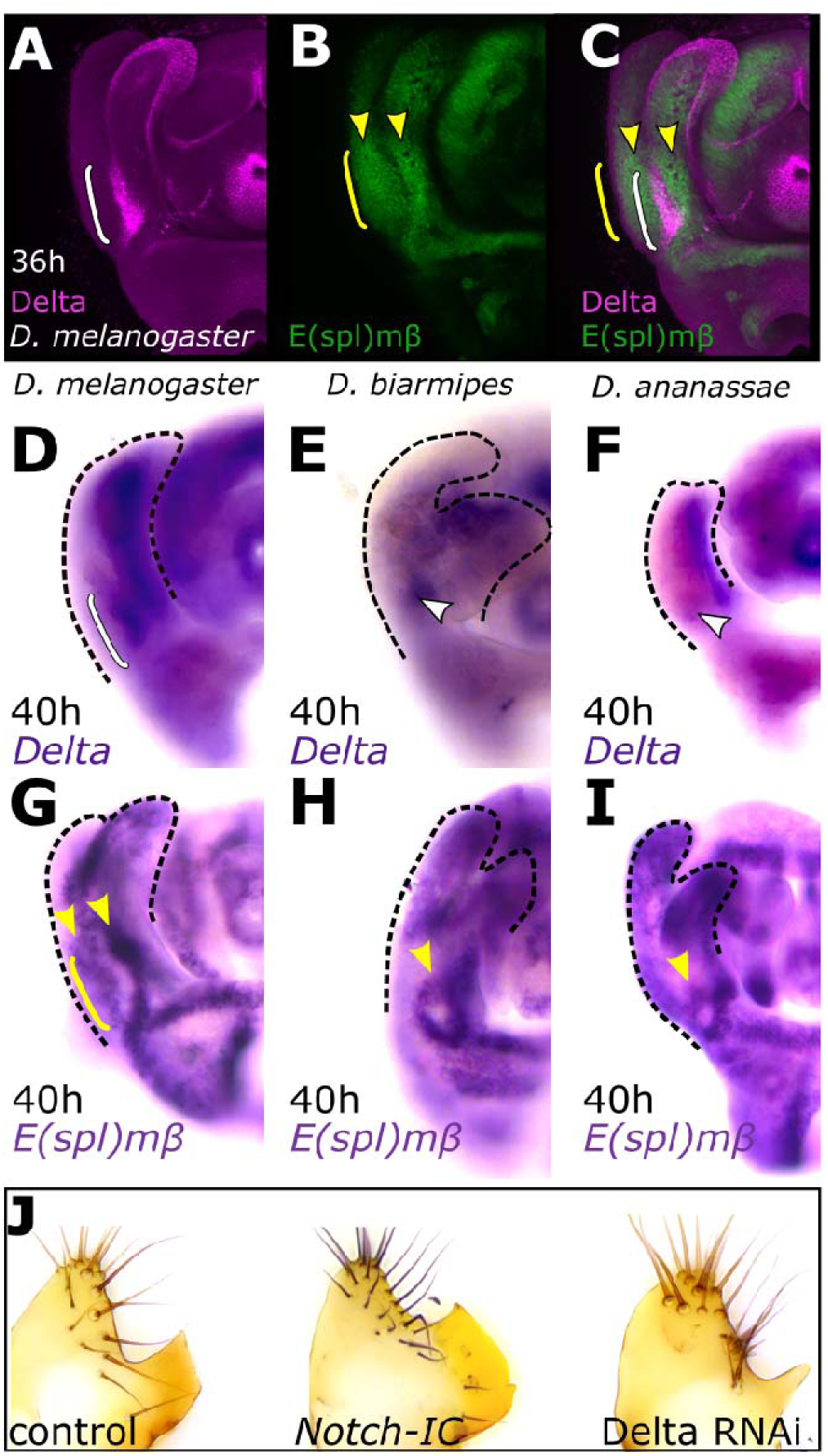
The spatially expanded expression of Delta is necessary for posterior lobe development in *D. melanogaster*. **(A)** Immunofluorescence of Delta protein expression (magenta). Delta expression is present adjacent to the developing posterior lobe at 36h APF in *D melanogaster* in a spatially expanded pattern (white bracket). **(B, C)** An E(spl)mβ-GFP reporter reveals Notch activity adjacent to Delta expressing cells (yellow arrowheads) and the cells of the developing posterior lobe (yellow bracket), and acts as a readout of Notch signaling. **(D-F)** Comparison of *Delta* mRNA expression between *D. melanogaster* **(D)** and the non-lobed species *D. biarmipes* **(E)** and *D. ananassae* **(F).** The expression of *Delta* expands in *D. melanogaster* (bracket), but is localized to a smaller region in non-lobed species (arrowheads, F and G). **(G-I)** Comparison of a canonical Notch target, *E(spl)m*β, mRNA expression between *D. melanogaster* **(G)**, *D. biarmipes* **(H)** and *D. ananassae* **(I).** The cells of the posterior lobe are Notch-responsive (yellow bracket). Two regions adjacent to either side of Delta expressing cells respond to Notch signaling (yellow arrowheads) **(G).** In non-lobed species, Notch activity is limited to a small circular pattern at the base of the lateral plate and clasper (yellow arrowhead in I and J). Represented images of B-J display half of a pupal genitalia. **(J)** Knockdown of Delta reduces the size of the posterior lobe (right), and over-activation of the Notch pathway increases the size of the posterior lobe (middle) compared to the control (left).

We next assessed whether the expanded pattern of posterior lobe-associated Delta is required for lobe development. A *Delta*-directed short hairpin RNA was driven by a genital-specific GAL4 driver of *Pox neuro* (*Poxn*) (42) which is active in a broad pattern in the genitalia, including the base of the lateral plate and claspers where *Delta* is also expressed (Fig. S2G) (22,42). Comparisons of adult phenotypes revealed that reduction of *Delta* expression interferes with posterior lobe development, resulting in smaller and defective posterior lobes (Fig. 2J and Fig. S2H, L-P). Importantly, reduction of *Delta* expression using this driver results in a pattern that resembles Delta expression in non-lobed species, suggesting that some aspect of the spatially expanded pattern of Delta is necessary for posterior lobe development (Fig. S2E, F and J). To further investigate the role of Delta-Notch in the development of the posterior lobe, we stimulated Notch pathway activity by expressing a constitutively active form of Notch (Notch intracellular domain) (43,44) under the control of the aforementioned *Poxn* driver. These animals developed a larger posterior lobe compared to the controls (Fig. 2J and Fig. S2H). This data showed that not only is the spatially expanded expression of *Delta* required for posterior lobe development, but that the size of the posterior lobe is sensitive to the amount of Notch signaling, suggesting that expansion of this pathway could have been an important evolutionary step in the origins of this structure.

To ascertain how the Notch pathway contributes to posterior lobe development, we assessed the spatial extent of its activation in the developing male genitalia. Delta is a transmembrane ligand of Notch, sending signals to adjacent cells (45,46). This cell-cell signaling is noteworthy because the expression of Delta expands between the lateral plate and clasper, which is adjacent to the developing posterior lobe. Thus, we expect the cells of the posterior lobe to be Notch responsive, receiving signals from the adjacent Delta pattern. To identify an appropriate readout of Notch activity, we tested the expression of canonical Notch targets of the *Enhancer of split* (*E(spl)*) complex by *in situ* hybridization (43). Throughout the entire genitalia, we found regions of bHLH repressor *E(spl)mβ* expression adjacent to regions of Delta expression, indicating that *E(spl)mβ* acts as an appropriate marker for Notch activity (Fig. 2G and Figs. S3-4). Notably, in *D. melanogaster*, *E(spl)mβ is expressed in* two patches of cells adjacent to each side of the expanded Delta pattern, including the posterior lobe developing from the lateral plate (Fig. 2G and Figs. S3-4). To directly compare *E(spl)mβ* with Delta, we constructed an *E(spl)mβ -*GFP transcriptional reporter transgene containing the proximal 1.4 kilobase (kb) of sequence adjacent to its promoter, a region known to recapitulate *E(spl)mβ* in other imaginal tissues (47). We observed that Delta and *E(spl)mβ*-GFP are indeed active in mutually exclusive regions and that the developing lobe showed apparent Notch pathway activation (Fig. 2B and C). To investigate Notch activity in non-lobed species, we tested the expression of *E(spl)mβ* in pupal genitalia of the non-lobed species *D. biarmipes* and *D. ananassae* by *in situ* hybridization. Similar to *D. melanogaster*, we found *E(spl)mβ* expression adjacent to Delta expressing cells*, but* the expression *of E(spl)mβ* was limited to a small ring-like pattern at the base of the lateral plates and claspers. The region of this ring-like pattern of *E(spl)mβ* mRNA appeared to anticorrelate with *Delta*’s spatially restricted expression patterns in these non-lobed species (Fig. 2H, I and Figs. S3-4). These observations indicate that 1) the cells of the posterior lobe are indeed Notch responsive and that 2) Notch/Delta signaling has a conserved ancestral pattern of downstream pathway activity during the development of genital structures.

### Partially redundant transcriptional enhancers regulate the lobe-associated expression of *Delta*

Considering the unique association of Delta’s expanded expression with lobe development, we sought to discover how this novel deployment occurred. To determine whether changes in *Delta* expression were encoded by *cis*-regulatory evolution, we first identified the enhancer sequences regulating *Delta* specifically in the lobe-forming region. To this end, we carried out a screen of ∼ 125 Kb of DNA at the *Delta* locus for pupal genital enhancers (Fig. 3). We utilized transgenic reporters from the Janelia GAL4 collection (48), which spanned ∼64 Kb of non-coding DNA upstream and intronic sequences of *Delta*. In addition, we cloned ∼95 Kb of non-coding DNA downstream of *Delta* into transgenic GFP (Green Fluorescent Protein) reporter constructs (Fig. 3A). Among these constructs, we identified two elements that recapitulate endogenous posterior lobe-associated expression of Delta (Fig. 3A-C). Enhancer1 is located ∼36 Kb downstream of the *Delta* transcription start site (Fig. 3A and B), and enhancer2 is ∼60 Kb downstream of the *Delta* coding transcription start site (Fig. 3A and C).

**Fig. 3.**
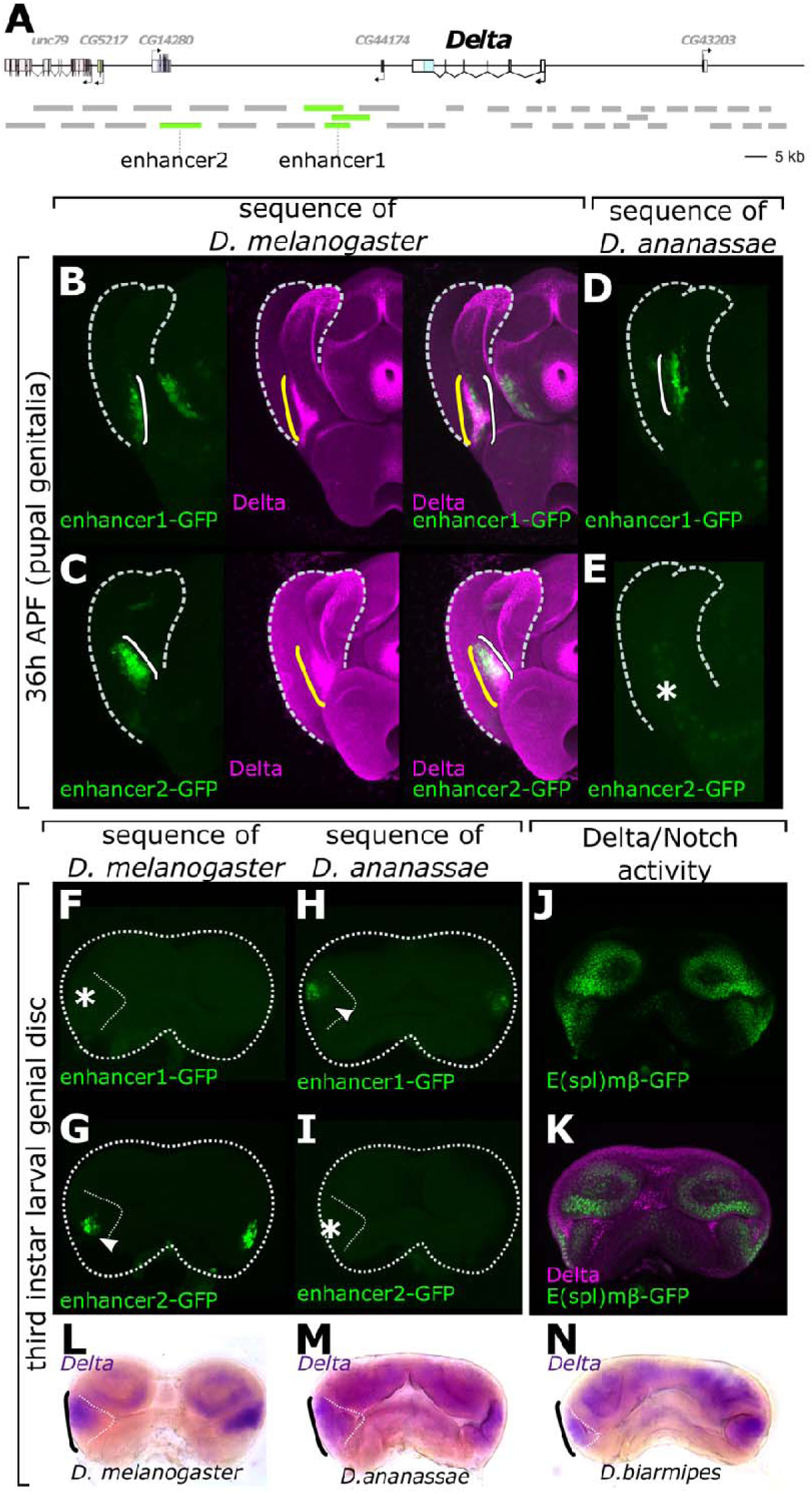
Two posterior lobe-associated enhancers of *Delta* are temporally stratified and have partially redundant spatial activities in the pupal genitalia. **(A)** schematic of the *Delta* locus, with two enhancer regions displayed by green bars downstream of *Delta.* The smaller green bar of enhancer1 marks the overlap of two larger fragments. Gray bars indicate tested regulatory regions in the locus. **(B, C)** Activity of the posterior-lobe associated enhancers of *Delta* (green, white bracket) relative to the endogenous expression of Delta (magenta, yellow bracket). Enhancer1 overlaps the full expansion of Delta **(B)** whereas enhancer2 recapitulates the more ventral portion of the pattern **(C). (D-E)** changes both in *cis* and *trans* contribute to the expanded expression of *Delta*. The orthologous region of enhancer1 from *D. ananassae* drives expression in an expanded pattern (bracket, **D**) similar to *D. melanogaster* (bracket, **B**) suggesting this element is functionally conserved. The orthologous segment of enhancer2 from *D. ananassae*, however, is not (or weakly) active in a *D. melanogaster* background (asterisks, **E**) suggesting changes to the element itself. **(F, G)** enhancer2 (arrow, G), but not enhancer1 (asterisk, F), is active in the larval genital imaginal disc. **(H, I)** the orthologous sequence of enhancer1 but not enhancer2 of *D. ananassae* drives larval disc activity in a *D. melanogaster* background. (**J, K**) Delta immunofluorescence and E(spl)mβ reporter activity reveal Notch activity in the larval genital disc (**J**) and Delta expression that recapitulates reporter activity in the larval disc (**K**). **(L-N)** *in situ* hybridization of *Delta* mRNA (black brackets) confirms *Delta* is expressed in the larval genital disc of *D. melanogaster* **(L)**, *D. ananassae* **(M)** and *D. biarmipes* **(N)**.

Having located the posterior lobe enhancers of *Delta*, we next investigated their functions and necessity in posterior lobe development. CRISPR/Cas9-mediated homology directed repair was employed to delete each enhancer individually. For each deletion, the enhancer region was replaced with a Ds-Red marker driven by a 3X-P3 promoter, which permitted the detection of integration events by eye fluorescence. To our surprise, neither enhancer deletion (2.4 kb Δenhancer1 or 5.5 kb Δenhancer2) reduced the size of the posterior lobe (Fig. S3A) or noticeably affected the expression of *Delta* (Fig. S5B). Notably, the minimal DNA sequences sufficient to drive reporter activity for enhancer1 and enhancer2 were ∼ 730 bp and ∼ 900 bp, respectively, and our deletion assays targeted a larger region to ensure the removal of potential nearby regulatory sequences important for the activity of these enhancers endogenously (Fig. S6). These results suggest that the two enhancers may act redundantly in regulating *Delta* near the posterior lobe.

### The Delta/Notch signaling center is active days before the posterior lobe forms

While the two enhancers of *Delta* have redundant activities and both overlap the endogenous expression of Delta, they exhibit slightly different spatial activities. In contrast to enhancer1, which fully recapitulates the endogenous expression pattern of *Delta* during posterior lobe development, enhancer2 is active in cells located more ventrally, between the lateral plate and clasper (Fig. 3B and C). Considering that expression patterns can change over developmental time, we were curious to know if the posterior lobe-associated enhancers of *Delta* showed differences in their timing. Thus, we investigated the relative developmental timing of each enhancer’s activation. To capture reporter activity more directly than GFP protein which persists for hours after expression, we performed *in situ hybridization* to detect *GFP* transcripts in the transgenic reporter lines (49)(Fig. S7A). This confirmed that the late activity of both enhancer1 and enhancer2 in the pupal genitalia represents active transcription rather than perdurance of GFP from an earlier stage. At earlier stages of pupal development, when the precursor of the posterior lobe has not yet differentiated, we detect stronger GFP mRNA signal from enhancer2 compared to enhancer1 (Fig. S7A). Considering that the posterior lobe develops during mid-pupal stages, the earliest pupal stages we had previously assessed were at 24h APF (hours after pupal formation). The reasoning behind this was that at 24h APF, the lateral plate which is the precursor of the posterior lobe has not fully separated from the clasper. Additionally, the developing genitalia of lobed and non-lobed species at this time point look quite similar, with the presumptive lateral plate and clasper still being fused together. Lastly, we were unable to detect obvious differences in Delta expression between lobed and non-lobed genitalia at 24h APF. However, the striking difference between enhancer1 and enhancer2 activity at 24h APF prompted us to assess even earlier pupal time points. To our surprise, enhancer2 was active throughout pupal development (Fig. 4A). To find the onset of enhancer2 activation, we tested its activity in a preceding stage of the *Drosophila* life cycle, the third instar larvae (L3), when the genital tissue is still an uneverted imaginal disc. Surprisingly, enhancer2 drove GFP expression in the male genital primordium of the imaginal disc in a bilaterally symmetric pattern (Fig. 3G) but GFP was undetected for the enhancer1-GFP reporter (Fig. 3F). We confirmed that the larval disc activity of enhancer2 recapitulates the endogenous expression of *Delta* by *in situ* hybridization and immunofluorescence (Fig. 3K,L and 4D, Fig. S8). Importantly, the *E(spl)mβ*-GFP reporter was active in mutually exclusive regions with Delta in the larval genital disc (Fig. 3J), suggesting Notch activity in response to the signaling center begins pre-pupal development. The early imaginal disc activity of Delta and enhancer2 was a particularly unexpected finding, as this stage occurs three days prior to the transition to the pupal stage, and about three and a half days before the initiation of posterior lobe development (50). Not surprisingly, this early activity is conserved between *D. melanogaster* and the non-lobed species *D. ananassae* and *D. biarmipes* (Fig. 3L-N, Fig. S8), suggesting that an early ancestral role of *Delta* exists that has been expanded upon and modified in *D. melanogaster* to produce a pattern required for the formation of the novel posterior lobe.

**Fig. 4.**
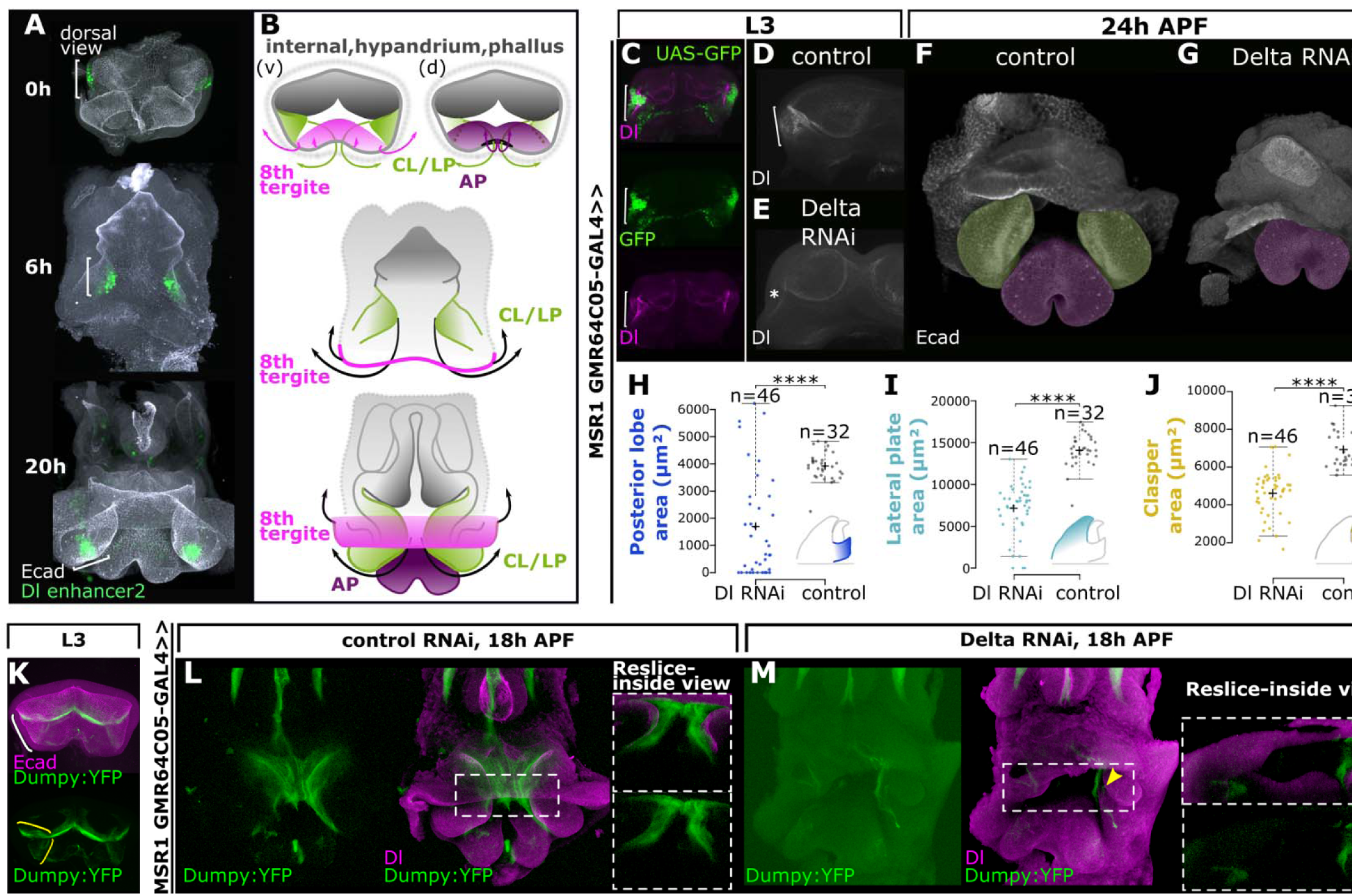
The posterior-lobe associated signaling center has an early ancestral role involved in genital disc eversion. **(A)** The activity of enhancer2 tracks with the cells of the genital disc that undergo evagination. At 0h APF, enhancer2 is active in the male genital primordium in a bilateral pattern. As the genital disc elongates at an early pupal stage (6h APF), enhancer2 tracks with the epithelial folds which will be a point of evagination. At 20h APF as the genitalia is mid-eversion, the signaling center marks a patch of cells between the prospective lateral plates and claspers. **(B)** cartoon description of panel A. The male genital primordium on the lateral sides of the L3 disc where Delta is active is denoted by green shading through development. The 8th tergite on the ventral side of the disc is colored in pink. This tissue everts along with the future clasper/lateral plate (CL/LP) and eventually fuses dorsally. The anal pate is colored in purple. Grey color marks tissue that will form the internal genitalia as well as the hypandrium, phallus, and branches. **(C)** Reporter activity of the MSR1-GMR64C05-GAL4 driver (green) in the L3 genital disc overlaps Delta expression (magenta) in the lateral folds of the male genital primordium. **(D, E)** Reduction of Delta by RNAi in the L3 genital disc (E, asterisks) compared to control (D, bracket). **(F, G)** Early knockdown of Delta in the L3 genital disc causes defects in genital eversion **(G)**. Prospective lateral plate/claspers (green false color) fail to evert in upon early Delta knockdown **(G)** compared to a control **(F).** The anal plate is false colored in purple, and future CL/LP in the control is false colored in green. **(H-J)** Reduction of Delta early in the L3 genital disc causes severe defects in the development of the posterior lobe **(H)**, lateral plate **(I)**, and clasper **(J)**. Asterisks denote significant differences (two tailed Student’s t-test, ****p<0.0001) **(K)** Dumpy is deposited within the L3 genital disc and covers the epithelial folds of the lateral male genital primordium. The lateral fold is marked by a white bracket (top) and Dumpy deposition is marked by yellow brackets (bottom). **(L, M)** Dumpy deposition is disrupted in genitalia with genital disc defects. Early knockdown of Delta with an L3 genital disc driver reduces and disrupts the organization of Dumpy tethers (compare G to F). Yellow arrowheads in panel M mark disrupted Dumpy localization in a Dl RNAi background.

Having identified an early role for enhancer2 in the larval imaginal disc, we reasoned that deletion of this enhancer would eliminate the early conserved activity of Delta and thus illuminate the ancestral role of this signaling center. However, as mentioned earlier, deletion of this enhancer did not affect Delta expression or the development of the posterior lobe. Furthermore, the early ancestral pattern of Delta was unaffected in the male genital primordium in Δenhancer2 animals (Fig. S5B). The lack of an obvious phenotype in this deletion line motivated us to further examine the *Delta* locus for additional enhancers, focusing on the larval genital disc. Utilizing the Janelia Gal4 collection (51), we identified seven additional upstream and intronic regions that drive larval disc activity (Fig. S10). While the expression domain of some of these reporters is quite broad, they all overlap Delta’s expression in the male genital primordium bilaterally on each side of the disc. Thus, it is likely that some of these enhancers work together or redundantly to regulate the early larval disc activity of Delta, suggesting that an early conserved role of Delta is controlled by a robust regulatory mechanism. Given that enhancer1 and enhancer2 drive strong pupal activity resembling Delta’s expression, we next examined the evolutionary history of these two enhancers.

### Changes within the *Delta* locus, as well as the *trans*-regulatory environment, contributed to the expansion of *Delta*

The spatial expansion of *Delta* in lobed species could be explained by two main mechanisms. First, mutations within the *cis*-regulatory elements may have led to *Delta*’s deployment in a larger population of cells in lobed species. Second, changes to *Delta*-regulating transcription factors may have caused this spatial expansion. To understand the evolutionary history of *Delta*’s posterior lobe-associated expansion and test these two possibilities, we examined the ability of the orthologous DNA fragments from the non-lobed species *D. ananassae* and *D. biarmipes* to drive GFP in a common *D. melanogaster trans*-landscape. Each DNA fragment was cloned into a GFP reporter construct and inserted into the same landing site in the genome of *D. melanogaster*. We found that the orthologous sequence of enhancer1 from each species is able to drive GFP in a pattern resembling *Delta*’s expanded expression (Fig. 3D and S7E). This recapitulation suggests that enhancer1 is functionally conserved and that changes within the *trans*-regulatory landscape have contributed to the expansion of *Delta*. In contrast to enhancer1, the orthologous sequence of enhancer2 from non-lobed species did not activate GFP in a *D. melanogaster* background, suggesting evolutionary modifications to this element (Fig. 3E and S4B-D).

Interestingly, during larval development, enhancer1 sequences from non-lobed species drove strong larval disc activity (Fig. 3H and S7E), unlike the *D. melanogaster* enhancer1. Enhancer2, on the other hand, drove little to no activity (Fig. 3I and S7E). These results suggest that while the genital imaginal disc activity of *Delta* is ancestral, the functionally conserved enhancer1 has lost an ancient role while the evolutionarily novel enhancer2 has taken over this conserved role. Collectively, comparative analysis of both enhancers indicated that a combination of *cis*- and *trans*-regulatory changes were responsible for the expansion of *Delta* to form the posterior lobe, and the two enhancers exhibit temporal shifts in the responsibility to drive the early conserved activity of Delta.

### Delta plays an ancestral role in the eversion of the genital disc

We hypothesized that *Delta* is responsible for an ancestral function because of the following considerations: 1) *Delta* was expressed in the imaginal genital discs of both lobed and non-lobed species, 2) *Delta*’s expression persisted in the developing pupal genitalia of non-lobed species, and 3) Notch signaling was active in regions adjacent to *Delta* expressing cells in the developing non-lobed genitalia. Furthermore, there is not a clear relationship between larval disc folds and pupal genital structures, and we were uncertain about which pupal structures the Delta expressing lateral regions of the L3 primordium corresponded to. Thus, we sought to investigate the ancestral function of *Delta* to understand the origin of this signaling center, and reasoned that tracing the lineage of *Delta* expressing cells would provide insight into its ancestral role. We performed a developmental time-course of enhancer2 activity from the genital imaginal disc through pupal development, staining with an antibody against E-cadherin to mark the apical surface of epithelial cells. Importantly, the genital imaginal disc starts inside out, with the apical surfaces facing each other and the basal lamina surrounding the outside of the disc. As the disc develops, this tissue turns partially inside out like a pillowcase, with external structures emerging from the posterior edge to form the external anal and genital structures, such as the lateral plates and claspers (52). At 0h APF (after pupal formation), enhancer2 is active in the male genital primordial folds on the lateral sides of the disc (Fig. 4A and B). The folds in which enhancer2 is active correspond to regions that begin to evert outward from the opening at the posterior stalk at early pupal stages (Fig. 4A and B). At around 20 h APF, the cells marked by enhancer2 have partially everted, which form the precursor of the lateral plate and clasper (Fig. 4A and B). These results demonstrated that *Delta*’s expression tracks with cells that undergo eversion to form the lateral plates and claspers, structures that are ancestral to the posterior lobe. Importantly, tracking the early larval disc expression of Delta with enhancer2 in *D. melanogaster* suggested that it is the same signaling center that persists through pupal development and later becomes precisely those Delta-positive cells adjacent to the posterior lobe as it forms.

To elucidate the ancestral role of *Delta*, we genetically perturbed the early expression of *Delta* in the larval imaginal disc. We utilized the publicly available Flylight Image Database (51) to select GAL4 drivers that are active in the lateral male genital primordium of the larval disc, and carried out a screen with the goal of knocking down *Delta* sufficiently early to specifically disrupt the early ancestral function. From this UAS-GAL4 *Delta*-RNAi screen, we successfully identified a GAL4 driver in the *MSR1* locus (GMR64C05-GAL4) that knocked down Delta in the lateral clusters of the male genital primordium without affecting *Delta*’s expression in other regions (Fig. 4C-E). As a result, 100% of the early *Delta* knockdown animals exhibited defects in genital eversion (Fig. 4G and S11C). At a pupal time point when the precursor of the lateral plates and claspers have everted in the control animals (Fig. 4F), the genitalia of *Delta* knockdown animals were still encapsulated within the tissue that should form the 8^th^ body segment (Fig. 4G). Interestingly, 32% of male adults completely lacked external genitalia, which we postulate is due to failure of eversion (Fig. S11A-B). Because RNAi is often incomplete and variable, *Delta* was not completely knocked down in some animals. Nonetheless, the adults that formed external genitalia had defects in either claspers, lateral plates, posterior lobes, or all three structures (Fig. 4H-J and S11D). These results indicate that Delta possessed an early function important for genital disc eversion, a complex process which is presumably necessary for genital development of all *Drosophila* species and predates the evolution of the posterior lobe.

### Delta is upstream of a vast apical extracellular matrix network

Our previous work on the cellular development of the posterior lobe revealed that the cells of the posterior lobe drastically increase in height to project from the developing lateral plate, and implicated an important role for the apical extracellular matrix (aECM) in shaping the posterior lobe cells (40). A gigantic protein of the aECM, Dumpy, covers the lateral plate of *D. melanogaster* in an expanded pattern compared to non-lobed species and is essential for proper posterior lobe development (40). It is speculated that Dumpy provides structural support as the intrinsic factors of the posterior lobe cells drive elongation, or that Dumpy tethers create a mechanical force to pull the cells of the posterior lobe (40). Given that Dumpy is a cellular effector molecule and likely downstream of the posterior lobe gene regulatory network, we were curious to know if there is a potential link between Delta and *dumpy*. Similar to *Delta*, *dumpy* expression is spatially expanded in *D. melanogaster* compared to non-lobed species, resembling the Notch responsive regions as the posterior lobe forms (40). Therefore, we examined whether Dumpy is downstream of Delta and whether it also has an early role in genital eversion. First, utilizing a line in which the Dumpy protein is endogenously tagged with a Yellow Fluorescent Protein (Dumpy:YFP) (54,55), we detected Dumpy deposition within the epithelial folds of the larval genital disc (Fig. 4K). Dumpy deposits within the genital disc clearly line the surface of the male genital primordium in the lateral regions, overlaying the Delta expressing epithelial cells (Fig. 4K and Fig. S12B). To test whether Dumpy has a role in the Delta-dependent genital disc eversion process, we examined whether Dumpy is disrupted under larval disc *Delta* RNAi conditions. In *Delta* RNAi animals with an eversion defect, Dumpy lost clear associations with the apical surface of the everting cells compared to the control which form concentrated deposits (Fig. 4L and M). Dumpy deposition in the larval genital disc was also decreased compared to controls (Fig. S12B), suggesting that the early activity of Delta is necessary for the presence of Dumpy at the surface of the male genital primordium. Together, these data suggest that Dumpy acts downstream of Delta during the eversion of the genital disc. In *Delta* RNAi conditions where the animals escaped the defective genital disc eversion phenotype, Dumpy localization was decreased on the prospective lobe-forming regions of the lateral plates (Fig. S14), suggesting that the novel role of Dumpy in posterior lobe development is also downstream of Delta.

## Discussion

Despite strong implications for the role of signaling pathways in the formation of many novel morphologies, efforts to understand how their pivotal roles were established have lagged far behind. Here, we identified a signal source, Delta, that is crucial to a novelty that formed on an intermediate timescale, permitting an opportune glance at its developmental and evolutionary past. Our comparative analysis of lobed and non-lobed species revealed that the Delta signal predated the posterior lobe novelty, suggesting roles preceding its evolution. The Delta signal initiates at a remarkably early developmental time, where it is required for the fundamental process of disc eversion, common to all species analyzed in this study. We placed this role in eversion upstream of the terminal effector Dumpy, which also participates directly in posterior lobe development. These findings suggest that morphological novelties that originate on macroevolutionary scales may evolve through elaborations of ancestral signaling centers whose beginnings may have been obscured by the passage of time. Because of their antiquity, the genetic architecture of these ancestral systems may be quite intricate and robust. We discuss below the implications of such “ancestral complexity” in understanding the origins of elaborate morphological structures.

The phenotypes we observed for *Delta* knockdown are some of the most dramatic posterior lobe defects we have found, and overexpression of the Notch intracellular domain is the only treatment currently known that dramatically increases lobe size to date. Considering the expanded expression of *Delta* near the developing lobe of *D. melanogaster,* this motivated our search for enhancers to trace Delta’s evolutionary history. While a combination of *cis* and *trans* differences accounts for the expanded expression we observed, the reporters notably allowed us to lineage trace the developmental trajectory of this signaling center back to the third instar genital disc. Prior to this study, our knowledge of the morphogenetic processes that occur during early genital disc development at pupal stages was sparse. Valuable studies in larval genital discs have identified genes and signaling pathways involved in anterior and posterior compartmentalization and patterning (56–58). While the eversion of the genital disc had previously been described (52), the molecular mechanisms governing this phenomenon remained largely uninvestigated. Our results highlight how novelties may depend upon signals initially deployed to pattern ancestral features at earlier stages of development. Work in the bat limb also hints at this possibility with the re-initiation of Shh signaling (13) specific to the bat wing after expression of Shh in the zone of polarizing activity has ceased. Such findings underline the importance of considering a wide range of developmental times that may bear upon evolving traits, rather than focusing solely on stages proximate to the developmental manifestation of the trait.

Our survey of *Delta* regulatory elements also revealed a redundant architecture that controls a broad domain of ligand expression that shifts over time. Redundant enhancers are quite common and are thought to foster robustness to environmental and genetic variation (59–61). The enhancer elements we found in the genital disc cover a wider spatial domain than was observed for the pupal enhancers (Fig S10). This suggests that the lobe-patterning signal center may have emerged from a portion of the total genital disc signaling center. Furthermore, our *Delta* knockdown experiments revealed a counterintuitive relationship between *Delta* expression and lobe patterning. When *Delta* RNAi is driven by the early *Delta* enhancer, the ventral expansion is completely ablated, but lobe morphology is unaffected (Fig. S9D-F). This suggests that the dorsal expansion (near the anal plate) represents the portion of the *Delta* signal relevant to the formation of this novelty. These results underscore the importance of fine-scale genetic manipulations to localize which portion of an expression pattern is most necessary to the developmental events under study.

Our previous work implicated a critical role of the aECM in the development of the posterior lobe (40). Here, we demonstrated that Dumpy deposition was affected when Delta was knocked down early in the larval genital disc, suggesting that Dumpy serves an early Delta-dependent role in disc eversion and a late role downstream of Delta in the posterior lobe network. A plausible model is that *dumpy* is regulated by Delta in the larval genital disc of both lobed and non-lobed species, carrying out an ancestral function of genital disc eversion. As *Delta* evolved an expanded expression pattern in *D. melanogaster*, the expression of *dumpy* may have expanded correspondingly. According to this model, a pre-existing target of Delta became involved in the development of the posterior lobe by becoming active in a broader expression domain (Fig. 5). Crucial to examining this model is identifying the regulatory elements that regulate *dumpy* in the larval genital disc as well as the posterior lobe. It is also possible, if not likely, that in addition to bringing along pre-existing downstream targets, the Notch pathway has also gained novel targets in lobed-species that contribute to posterior lobe development.

**Fig. 5.**
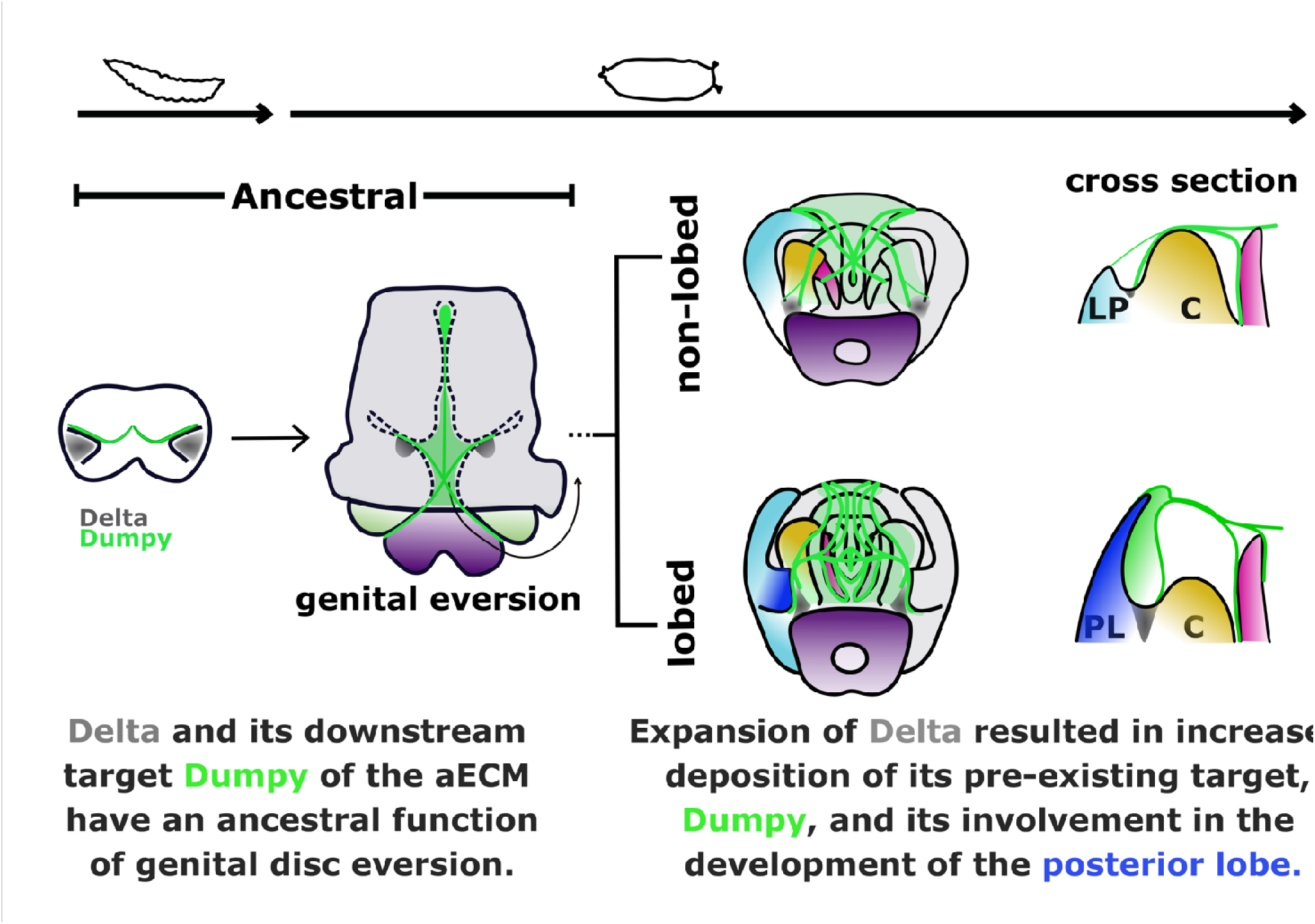
A Notch signal required for a morphological novelty in Drosophila has antecedent functions in genital disc eversion. (From left to right) A Delta/Notch signaling center (grey) is active in the larval genital disc with an ancestral function of genital disc eversion, a process that is conserved among *Drosophila* species. Dumpy (green lines), a component of the aECM, is associated with antecedent roles in genital disc eversion in a Delta-dependent manner and may be a pre-existing target of Delta. The Delta/Notch signaling center persists through pupal development in non-lobed species, where it carries out potential ancestral roles in later pupal development, such as lateral plate and clasper development through Dumpy connections on the epithelia of these structures. In lobe-forming species, the ancestral pattern at the base of the lateral plate/clasper precursor (shaded green) later spatially expands along the two structures through *trans* regulatory changes (lateral plate in light blue, clasper in yellow) and form the novel posterior lobe (royal blue). This model suggests that as the expanded expression of Delta evolved, its pre-existing downstream target, Dumpy, gained an additional activity involved in posterior development.

Our previous work on the evolution of the posterior lobe had shown that an ancestral network of the embryonic posterior spiracle was co-opted in the genitalia (22). The connection between the Delta signaling center and co-opted posterior spiracle network however is yet to be explored. An interesting question is whether the expansion of Delta is downstream of the co-opted network or vice-versa. Our preliminary data suggests that the expansion of Delta during pupal stages is downstream of Pox Neuro (Poxn), a key component of the co-opted network (Fig. S15). In a Poxn mutant, the ancestral (not-expanded) pattern of Delta remains unaffected, however, this pattern does not expand. This is in agreement with our data showing that Delta has a much earlier onset of expression than Poxn. Thus, it is possible that the derived, expanded expression of Delta is mostly attributed to *trans*-regulatory changes that occurred as a result of the co-opted spiracle network, and the co-option of the spiracle network may have ignited the redeployment of transcriptional regulators rather than terminal effectors.

The complexity of ancestral systems has been an important concept in developmental evolution. As we learned from sequencing the human genome (62) and an increasing array of basally branching organisms (63–65), the field realized that most transcription factors and signaling pathways are conserved within animals and beyond rather than each clade having an abundance of lineage specific regulatory genes and gene families. Our work here causes us to appreciate how ancestrally complex signaling centers may apply to newly formed novelties. These centers may show faint signs of relation to signals deployed much earlier in development. The regulatory architecture of their participating loci may show unexpected intricacies, such as redundancy that resulted from the evolutionary refinement of their ancestral roles. Our efforts to resolve these faint connections and complex developmental systems are crucial to developing a sophisticated understanding of what would otherwise appear to have arisen through inexplicable events.

## Materials and Methods

### *Drosophila* strains and husbandry

Fly stocks were reared at room temperature on standard cornmeal agar media. For RNAi experiments, flies were reared at 29 °C. The *Drosophila melanogaster* line used in this study is mutant for the *yellow* and *white* genes and was isogenized for eight generations (*y^1^w^1^*, Bloomington Stock Center #1495). Outgroup species that lack a posterior lobe *(Drosophila ananassae #*0000-1005.01, *Drosophila biarmipes #*0000-1028.01) were obtained from the University of California, San Diego Drosophila Stock Center (now called The National Drosophila Species Stock Center at Cornell University). All stocks are listed in Table S1

### Transgenic constructs

To generate GFP reporter flies, regulatory elements were PCR amplified using primers listed in Table x, and cloned into a vector containing GFP and a minimal hsp70 promoter (pS3aG) (66). Primers were designed using the GenePalette software tool (67). *AscI* and *SbfI* restriction sites were added to the primer sequences (Integrated DNA Technologies) to insert the amplified region into the multiple cloning site of the vector. Regions of interest were amplified from genomic DNA prepared by the DNeasy Blood & Tissue Kit (QIAGEN). *D. melanogaster* transformant lines were created by phiC31 mediated site specific recombination into the 68A4 “attP2” landing site on the third chromosome(68) or the 51D site on the second chromosome(69) by Rainbow Transgenics. For each GFP reporter, 2-5 independent insertion lines were analyzed.

### Larval and pupal genital sample preparation

Pupal genital samples were prepared for in situ hybridization according to Glassford et al. 2015 (22,70). In short, to standardize aging, male white pre-pupae were incubated at 25°C until ready for dissection. Pupae were cut in half in cold PBS, fat bodies were flushed out, and pupal case was removed. Samples younger than 20 hours APF (after pupal formation) and larval samples were cut in half and turned inside out at the posterior end to remove fat bodies to prevent damaging or dislodging the delicate early genitalia. All samples were fixed in PBS with 0.1% Triton-X and 4% paraformaldehyde (PBT-fix) for 30 min at room temperature. Samples containing fluorescent labels or being prepared for immunostaining were then washed twice in PBT. Samples to be used for in situ hybridization were rinsed twice in methanol and twice in ethanol, and stored at −20°C in 100% ethanol.

### Generation of CRISPR/Cas9 mutants

Deletion of the enhancers of *Delta* was accomplished by CRISPR/Cas9 homology directed repair using gRNA targets flanking the enhancer boundaries (Table S2). For deletion of enhancer-1, *D. melanogaster* embryos were injected by Rainbow Transgenics with a mixture of 250 ng/μL nos-Cas9 vector, 100 ng/μL of each gRNA vector (pCFD3-dU63gRNA), and 500 ng/μL donor plasmid. The donor plasmids for homology directed repair contain a *3XP3::DsRed* cassette flanked by approximately 1kb of genomic DNA. Transformants were identified by the expression of DsRed in the eyes of the progeny of the injected flies. Embryo injections to delete the early enhancer of *Delta* were performed in house. A mixture of 200 ng/μL per gRNA and 500 ng/μL donor vector was injected into nos-Cas9 expressing embryos (Bloomington #78781). gRNAs were generated by *in vitro* transcription. Briefly, a double stranded DNA template containing a T7 promoter was amplified by PCR. In vitro transcription was then carried out using the MEGAscript T7 Transcription kit (Invitrogen).

### In situ hybridization

To detect mRNA localization, in situ hybridization was performed following the protocol described in Rebeiz et al., 2009. Modifications were made according to Glassford et al., 2015 to utilize the InsituPro VSi robot (Intavis Bioanalytical Instruments)(70). Briefly, fixed samples were washed in methanol, rehydrated in PBT (PBS with 0.1% Triton-X), fixed in PBT-fix, and incubated in hybridization buffer for 1 hour at 65°C. Prehybridized samples were then incubated with digoxygenin riboprobes (primers for amplifying mRNA probes listed in Table X) for 16 hours at 65°C, and subsequently washed in hybridization buffer followed by PBT washes to remove unbound riboprobes. To reduce background noise, samples were blocked in PBT with 1% bovine serum albumin for 2 hours. Blocked samples were then incubated with anti-digoxigenin antibody Fab fragments conjugated to alkaline phosphatase (Roche Diagnostics) diluted in PBT at 1:6000 overnight at 4°C. After several PBT washes, alkaline phosphatase color reactions were performed by incubating samples in nitro-blue tetrazolium chloride and 5-bromo-4-chloro-3’-indolyphosphate p-toluidine salt (NBT/BCIP) (Promega) and monitored under a dissecting microscope. Color reactions were stopped by PBT washes when purple stain was detected. Samples were mounted on a glass slide coated with Poly-L-Lysine in an 80% glycerol 0.1 M Tris-HCL (pH 8.0) solution.

### Immunostaining

To detect the expression patterns of proteins, genital samples removed from the pupal membrane were incubated in primary antibody diluted in PBT at 4°C overnight. The following primary antibodies were used: monoclonal mouse anti-Delta 1:100 (DSHB, #C594.9B0s), polyclonal goat anti-Delta 1:100 (Santa Cruz Biotechnology, Inc.), rat anti-Ecadherin 1:100 (DSHB, #DCAD2). To remove unbound primary antibody, the samples were washed in PBT 3-5 times over the course of an hour. To detect bound primary antibody, the samples were subsequently incubated in a fluorescent-dye conjugated secondary antibody diluted in PBT and incubated either at 4°C overnight or at room temperature for 4 hours. The following secondary antibodies were used: donkey anti-mouse Alexa 488 1:400 (A21202, Thermo Fisher Scientific), donkey anti-goat Cy2 1:400 (705-225-147, Jackson ImmunoResearch), and donkey anti-rat Alexa 594 1:400 (A21209, Invitrogen). Samples were then washed in PBT, incubated in 50% PBT and 50% glycerol solution, and mounted on a glass slide in an 80% glycerol 0.1 M Tris-HCL (pH 8.0) solution. Mounting wells were made with a single layer of double sticky tape on the glass slides. To avoid rotation of the sample during mounting, the wells were coated with Poly-L-Lysine solution. Glass cover slips were placed on the samples to seal the wells.

### Microscopy and image analysis

Cuticles of adult genitalia and stained in situ hybridization samples were imaged on a Leica DM2000 with a Leica DFC540 camera at 20x magnification. Fluorescently labeled samples were imaged using an Olympus Fluoview 1000 or a Leica TCS SP8 confocal microscope at 40x magnification. Larval samples were imaged at 63x magnification where indicated. Images were processed with Fiji(71).

## Acknowledgments

The authors thank members of the M.R. laboratory for their comments and discussion on the manuscript. We thank Gerard Campbell for providing fly stocks and thoughtful discussions. We thank Werner Boll, Markus Noll, the Bloomington stock center, and Drosophila Genomics and Genetic Resources for fly stocks. We thank Charles Elliot for early preliminary studies that motivated this study.

## Author contributions

Conceptualization: DNS, MR

Methodology: DNS, MR

Investigation: DNS, WJG, GR SJS, MR

Resources: DNS, SJS

Visualization: DNS, MR

Formal Analysis: DNS, MR

Supervision: MR

Funding Acquisition: MR

Writing—original draft: DNS, MR

Writing—review & editing: DNS, WJG, GR, SJS, MR

## Data and materials availability

All data are available in the main text or the supplementary materials.

