## Supplementary material for "A Notch signal required for a morphological novelty in *Drosophila* has antecedent functions in genital disc eversion": supplmental data

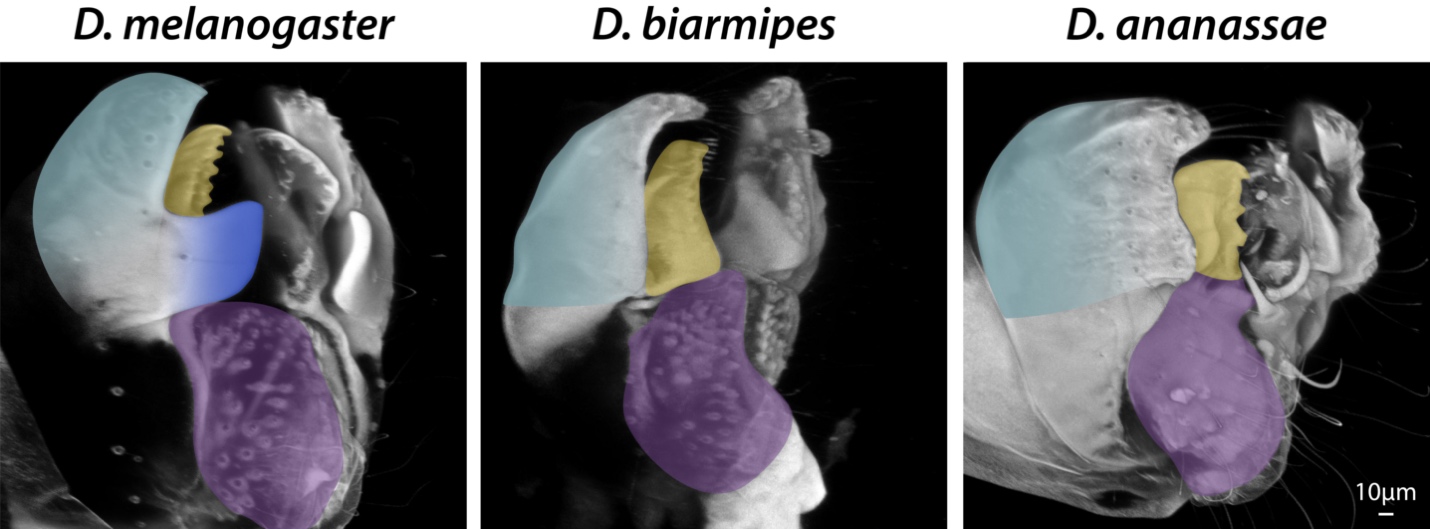

Fig. S1.

The posterior lobe is a novel morphological outgrowth of the lateral plate. Autofluorescent confocal images of *D. melanogaster*, *D. biarmipes,* and *D. ananassae* showing the anal plates (purple), clasper (yellow), lateral plate (light blue) and posterior lobe (dark blue). Images were rotated in 3D in the MorphoGraphX software package. Note that darkly colored cuticle (e.g. adjacent to the lateral plate and anal plates in *D. melanogaster* and *D. biarmipes*) does not autofluoresce despite the presence of cuticle.

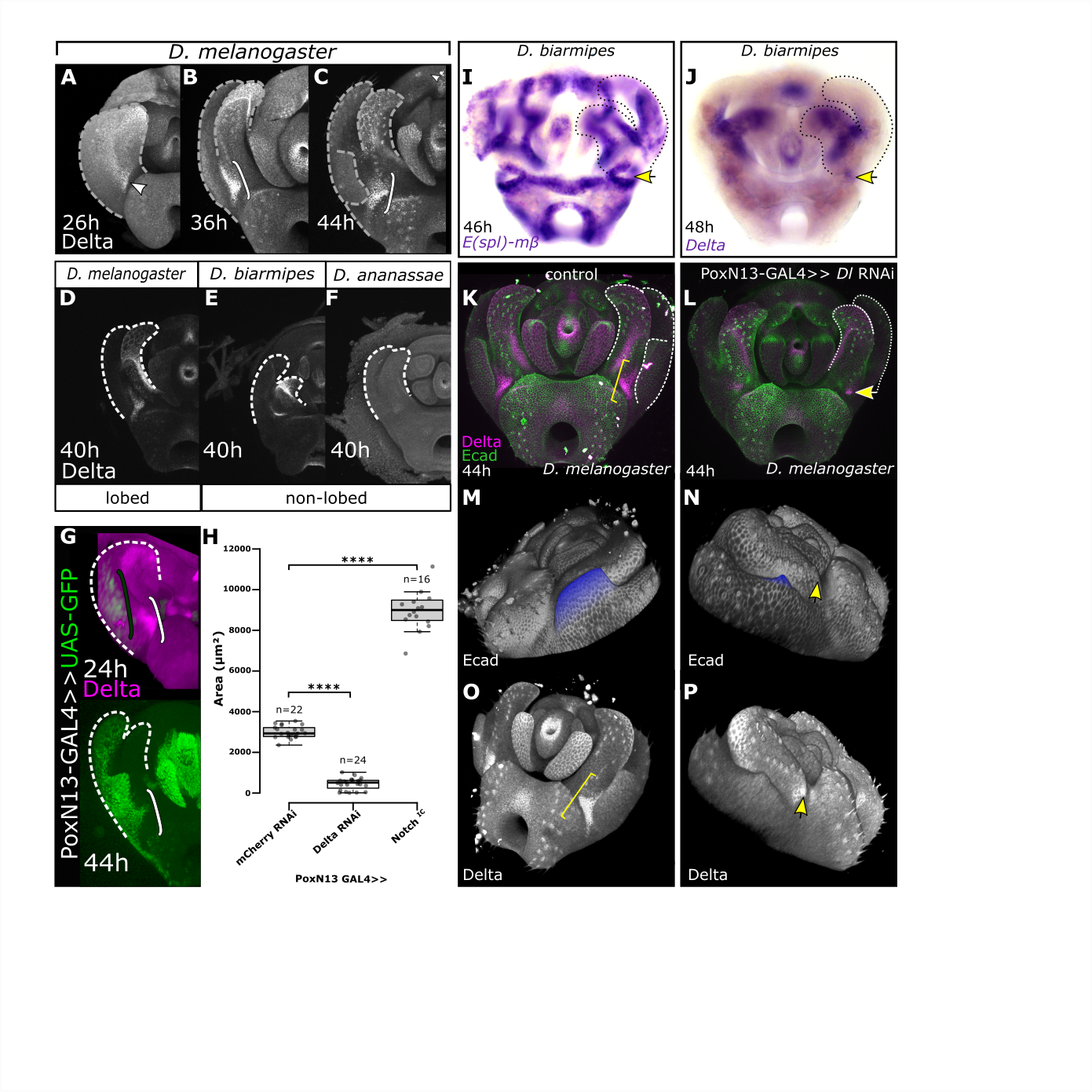

Fig. S2.

**The expanded expression of Delta is unique to the lobed *D. melanogaster* and is necessary for posterior lobe development.** (A-C) Immunofluorescent time-course of Delta expression. At 26h APF prior to the initiation of posterior lobe development the lateral plate and clasper are fused, and Delta is expressed at the dorsal base of the lateral plate/ clasper (arrowhead, A). As the lobe initiates its development at 36h APF, the expression of Delta spatially expands adjacent to the developing posterior lobe (bracket, B), and at 44h APF when the lobe has formed, the lobe-associated expression of Delta retracts (bracket, C). (D-E) Pupal genitalia of *D. melanogaster*, *D. biarmipes*, and *D. ananassae* stained with a cross-reactive polyclonal antibody against Delta. Expanded lobe-associated expression is marked with a bracket (D), and the ancestral pattern is indicated by arrowheads (E, F). (G) Reporter activity of the PoxN13-GAL4 driver (green) at 24h and 44h APF. At 24h APF while the driver is active in the presumptive lateral plate (bracket), its activity does not overlap Delta (magenta, white arrowhead). (H) Quantification of Delta RNAi and constitutive activation of the Notch pathway using the PoxN13- GAL4 driver. Asterisks denote significant differences (two tailed Student’s t-test, ****p<0.0001). Whiskers extend 1.5 times the interquartile range above and below Q1-Q3. (I, J) Spatial activity of Delta and Notch responsive regions in the non-lobed species *D. biarmipes*. *E(spl)mβ* mRNA is expressed in a tight circle at the base of the lateral plate and clasper in *D. biarmipes* (I, yellow arrow)), surrounding the restricted Delta expression pattern in this region (J, yellow arrow) which resembles the Delta knockdown pattern in *D. melanogaster* (compare J and L). (K-P) Knockdown of Delta with the PoxN13-GAL4 driver reduces the expression of Delta (compare white arrowhead in L to bracket in K), resembling its ancestral expression pattern (J). (M, N) 3D rendered confocal images of panels K and L depicting E-cadherin staining, rotated to display a wildtype posterior lobe in the control (M), and a reduced posterior lobe (N). Posterior lobes are false colored in royal blue. (P, O) 3D rendered confocal images of panels K and L depicting Delta staining, , rotated to display a wildtype posterior lobe in the control and the expanded Delta pattern (O, yellow bracket), and a reduced posterior lobe corresponding to a spatial reduction in Delta expression(P, yellow arrow).

Fig. S3.

**The expression of *Delta* and *E(spl)mβ* are anticorrelated.** (A-L) *In situ* hybridization of *Delta* mRNA in *D. melanogaster*, *D. biarmipes*, and *D. ananassae.* In the lobe forming species, *D. melanogaster,* Delta expression expands at the base of the lateral plate and clasper during lobe development. In the two non-lobed species, *D. biarmipes* and *D. ananassae*, however, *Delta* expression does not spatially expand (white arrowheads). (M-X) *In situ* hybridization of mRNA in *D. melanogaster*, *D. biarmipes*, and *D. ananassae* *E(spl)mβ* mRNA in *D. melanogaster*, *D. biarmipes*, and *D. ananassae.* In *D. melanogaster*, the Notch target, *E(spl)mβ,* is expressed in two patches adjacent to the Delta expressing region (yellow brackets). In *D. biarmipes* and *D. ananassae,* Notch activity is limited to a small circular pattern at the base of the lateral plate and clasper and does not extend to the lateral plate.

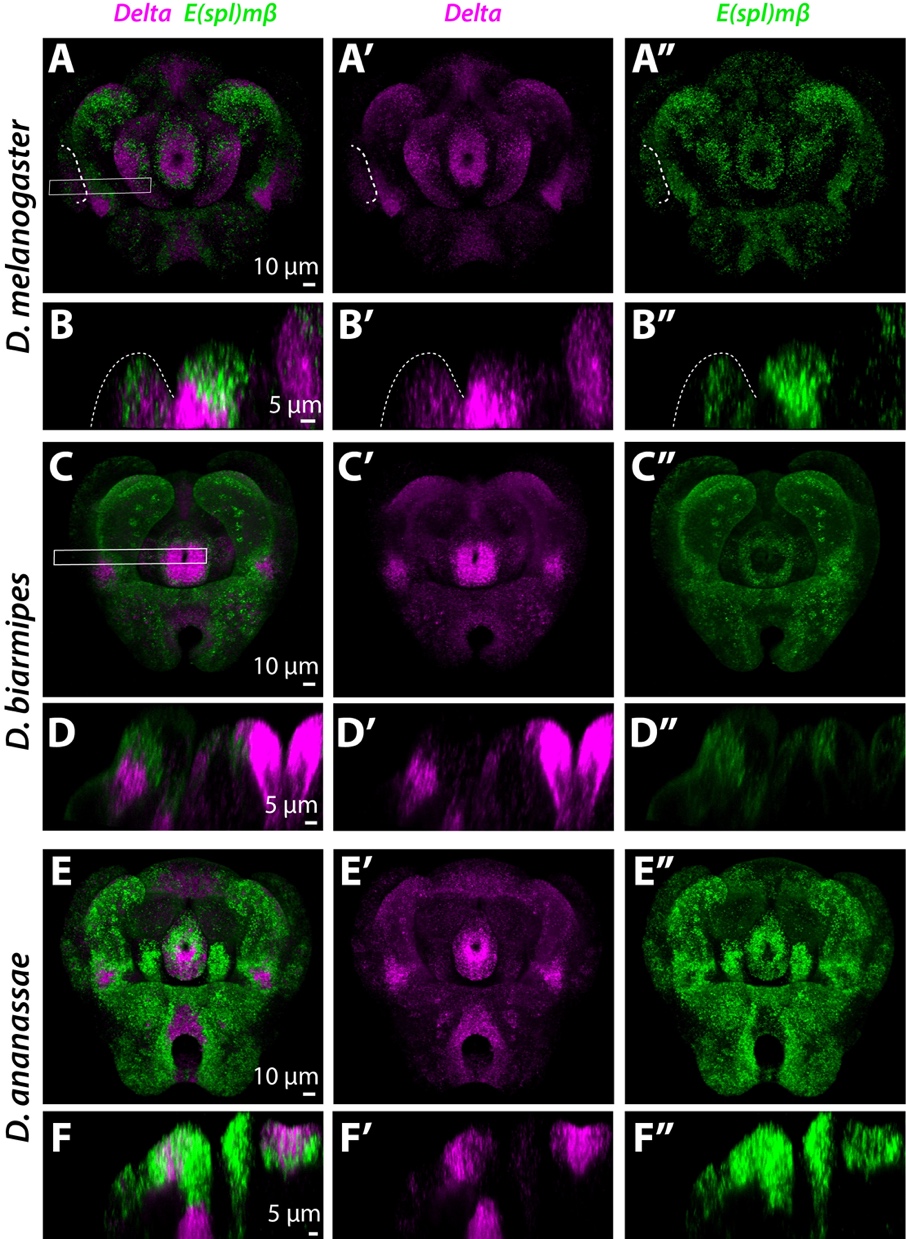

Fig. S4.

Three dimensional anticorrelation of *Delta* and *E(spl)mβ* expression. HCR was used to visualize *Delta* (purple) and *E(spl)mβ* (green) expression patterns in 40h APF of *D. melanogaster*, *D. biarmipes*, and *D. ananassae* genitalia. (A-A’’) In *D. melanogaster*, *Delta* and *E(spl)mβ* are expressed in the posterior lobe (dashed line) and the clasper (medial of the dashed line). (B-B’’) A cross-section highlights that *Delta* and *E(spl)mβ* expression appear to be non-overlapping. Note that *Delta* mRNA is detected in cells of the developing posterior lobe (B’), but we fail to detect protein expression in these cells (Fig. S2A). (C-C’’) In *D. biarmipes*, *Delta* and *E(spl)mβ* are expressed in the clasper (medial of the dashed line) but not the lateral plate (dashed line). (D-D’’) A cross-section highlights that *Delta* and *E(spl)mβ* expression appear to be non-overlapping. (E-E’’) In *D. ananassae*, *Delta* is expressed in the clasper (medial of the dashed line) but not the lateral plate (dashed line), while *E(spl)mβ* is expressed in both tissues. (F-F’’) A cross-section highlights that *Delta* and *E(spl)mβ* expression appear to be non-overlapping. White boxes outline the region used to make a cross-section stack.

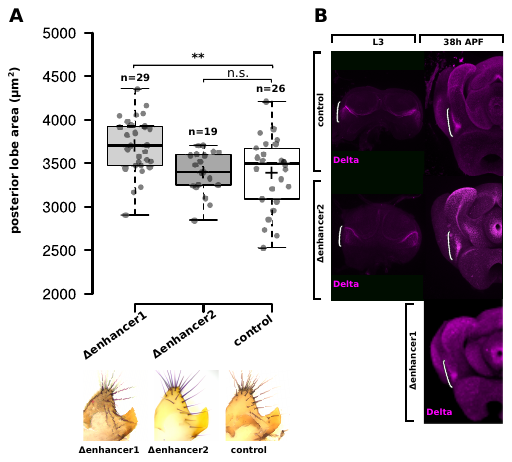

Fig. S5.

**Neither deletion of enhancer1 nor enhancer2 reduces the size of the posterior lobe.** (A) Quantification of each enhancer deletion. Asterisks denote significant differences (two tailed Student’s t-test, **p<0.01, n.s. denotes not significant). (B) Delta expression in L3 genital disc and 38h APF pupal genitalia of control (top) compared to ∆enhancer2 (middle) indicates Delta does not display qualitative difference in the absence of enhancer2 which drives both larval and pupal activities. In the absence of the pupal enhancer1, Delta is still expressed in an expanded pattern at 38h APF (bottom).

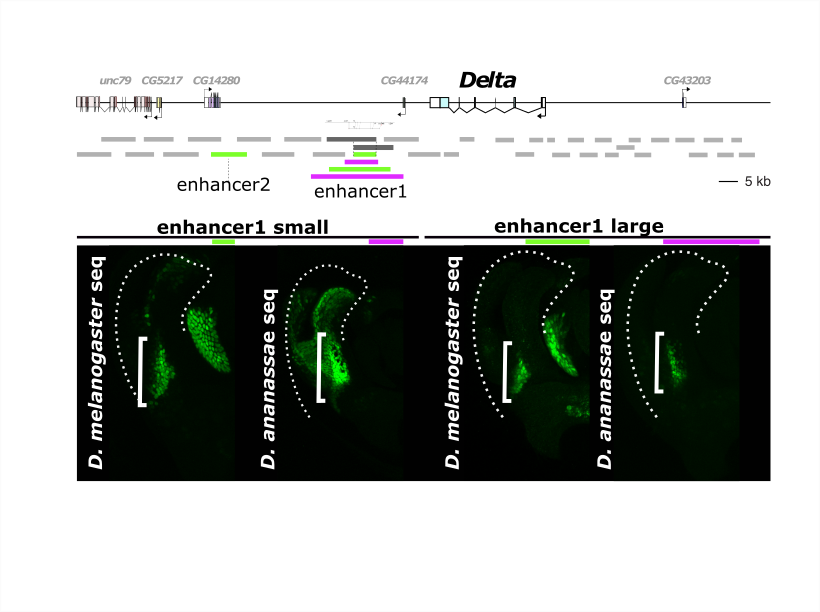

Fig. S6.

**Subdivision of *Delta* enhancer1.** (top) Map of the *Delta* locus, showing the position of large (green) and small (magenta) subdivisions of enhancer1. (bottom) GFP reporter images at 36hAPF of the large and small versions of the *D. melanogaster* and *D. ananassae* enhancer1 versions.

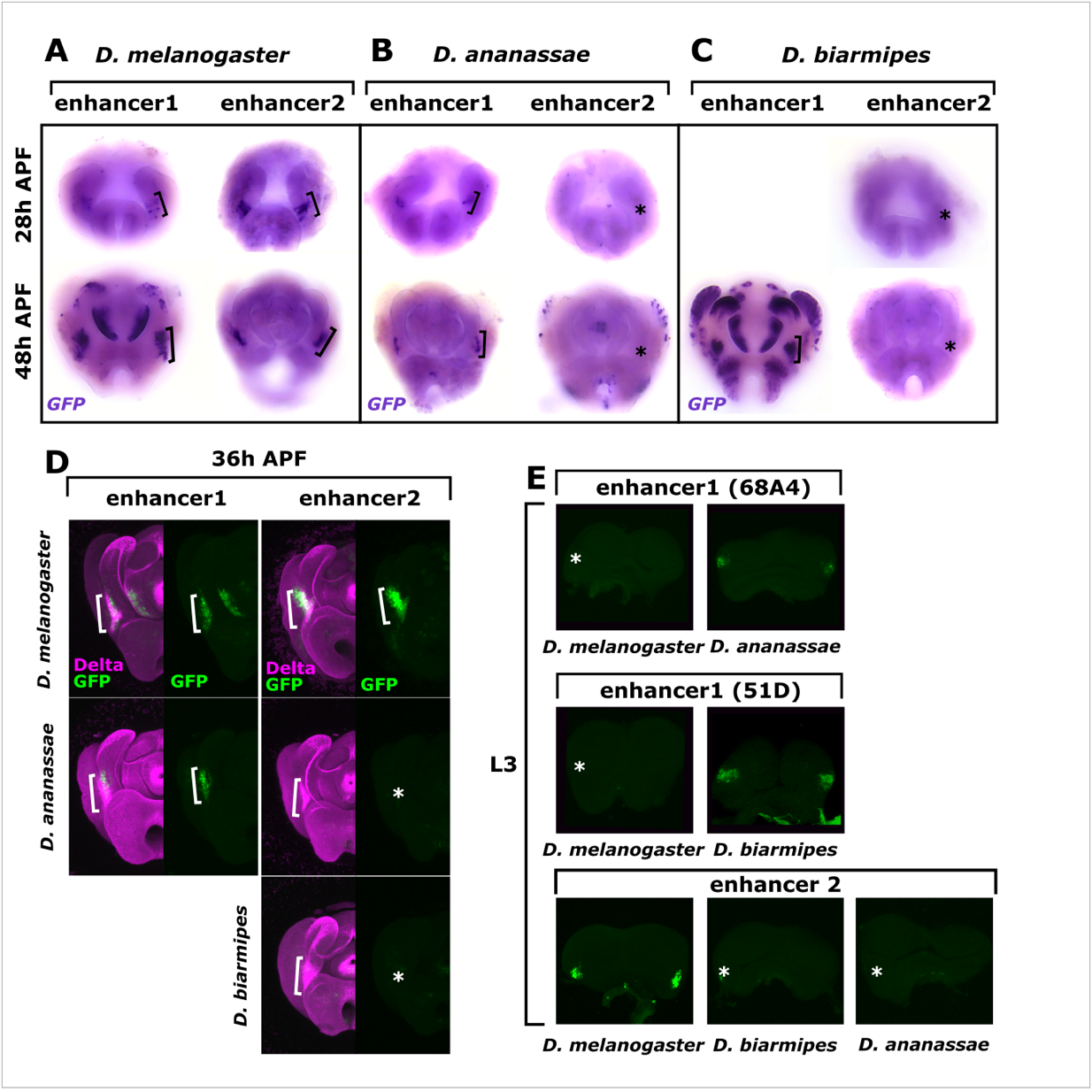

Fig. S7.

**Temporal and spatial activity of Delta enhancer1 and enhancer 2.** (A-C) GFP *in situ* hybridization of the lobe-associated reporters of Delta. The lobe-associated enhancers of Delta have differences in their spatial and temporal activities in *D. melanogaster* (A). In D. melanogaster, enhancer2 drives stronger activity at an early pupal stage (28h APF). As the lobe has developed at 48h APF, both enhancers still drive activity, albeit enhancer1 in a broader spatial pattern compared to the more ventrally active enhancer2. (B and C) The orthologous sequences of enhancer1 from *D. ananassae* (B) and *D. biarmipes* (C) drive pupal expression in a *D. melanogaster* background, suggesting the element is functionally conserved. However, the orthologous region of enhancer2 from these species fails to drive activity in the pupae (asterisks), suggesting that enhancer2 has been modified. (D) Comparison of orthologous reporters by GFP fluorescence in the developing pupal genitalia at 36h APF. Samples were stained against a monoclonal antibody against Delta. (E) Comparison of orthologous reporters by GFP fluorescence in the L3 genital disc. 68A4 and 51D denote landing sites in the D. melanogaster genome.

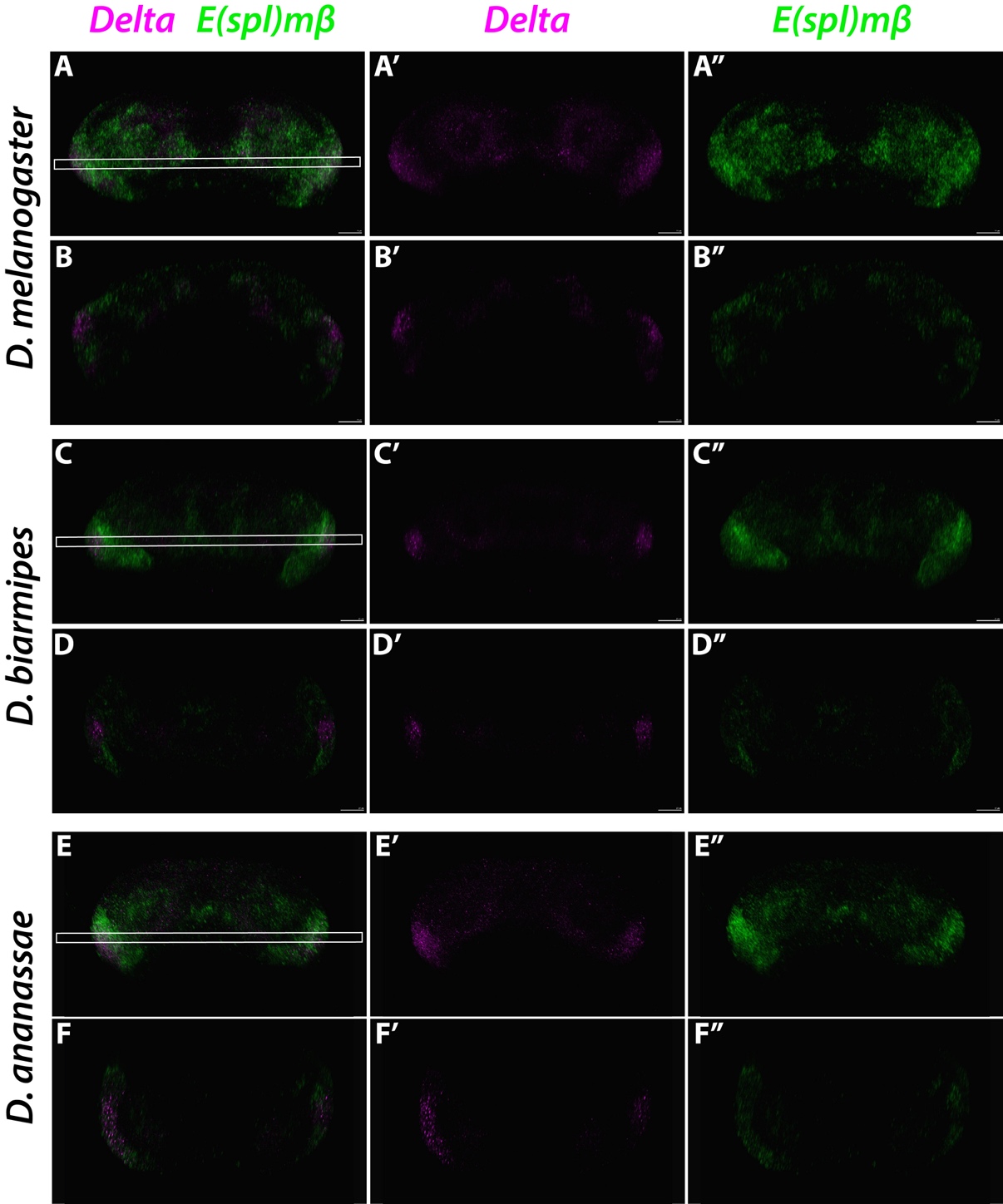

Fig. S8.

**Conservation of Delta expression in the L3 genital disc.**

HCR was used to visualize Delta (purple) and mBeta (green) expression patterns on L3 genital discs of *D. melanogaster*, *D. biarmipes*, and *D. ananassae*. (A-A’’) In *D. melanogaster*, *Delta* and *E(spl)mβ* are expressed in the lateral and medial regions of the L3 imaginal disc. (B-B’’) A cross-section highlights that *Delta* and *E(spl)mβ* expression appear to be non-overlapping. (C-C’’) In *D. biarmipes*, *Delta* and *E(spl)mβ* are expressed in the lateral regions of the L3 imaginal disc. (D-D’’) A cross-section highlights that *Delta* and *E(spl)mβ* expression appear to be non-overlapping. (E-E’’) In *D. ananassae*, *Delta* and *E(spl)mβ* are expressed in the lateral regions of the L3 imaginal disc. (F-F’’) A cross-section highlights that *Delta* and *E(spl)mβ* expression appear to be non-overlapping. White boxes outline the region used to make a cross-section stack.

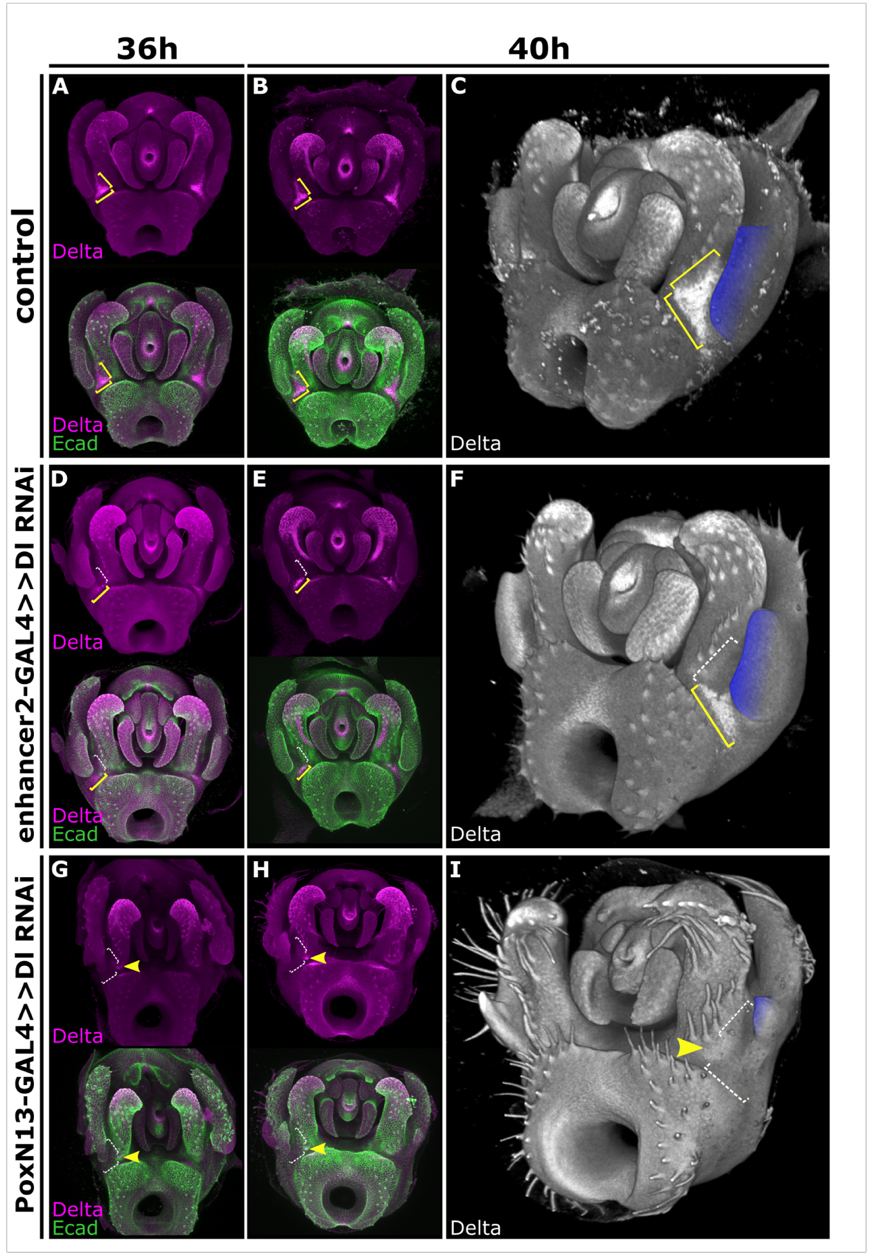

**Fig. S9.**

**Dissecting the expanded pattern of Delta to its functional sub-patterns.** (A-B) WT Delta expression at two timepoints by immunofluorescence. The expanded expression of Delta can be broken down into a ventral and dorsal extension (yellow brackets). Dorsal extension is towards the anal plate. Bottom panels show merges with E-cadherin antibody to visualize the apical surface of the cells. (C) 3D rendering of a 40h APF pupa stained with Delta (yellow brackets). The developing posterior lobe is false colored in blue. (B-F) RNAi knockdown of *Delta* using a GAL4 line driven by the early enhancer2 of Delta affects the ventral extension of Delta (white bracket indicates loss of ventral extension) but not the dorsal extension (yellow bracket). Posterior lobe development is not noticeably affected in these animals. (G-I) RNAi knockdown of *Delta* driven by PoxN-GAL4 affects both the dorsal and ventral expansions of Delta (white brackets) and causes a severe posterior lobe defect. Yellow arrowheads point to the remaining Delta activity at the base of the clasper.

Fig. S10.

Highly redundant regulatory elements overlap the larval disc activity of Delta. (A) schematic of the Delta locus, displaying the relative region of D. melanogaster regulatory elements driving larval disc activity. Green bar indicates activity in both L3 and pupal stages, white bar indicates pupal activity only, and grey bars indicate L3 activity all in D. melanogaster. (B-I) Confocal images of reporters in the male L3 genital disc (green) overlapping Delta expression (magenta) in the male genital primordium 51 located laterally on the disc (brackets). Images correspond to the order of annotated elements in panel A from left to right. Represented images are half of the L3 disc. confocal images were taken at 63x magnification.

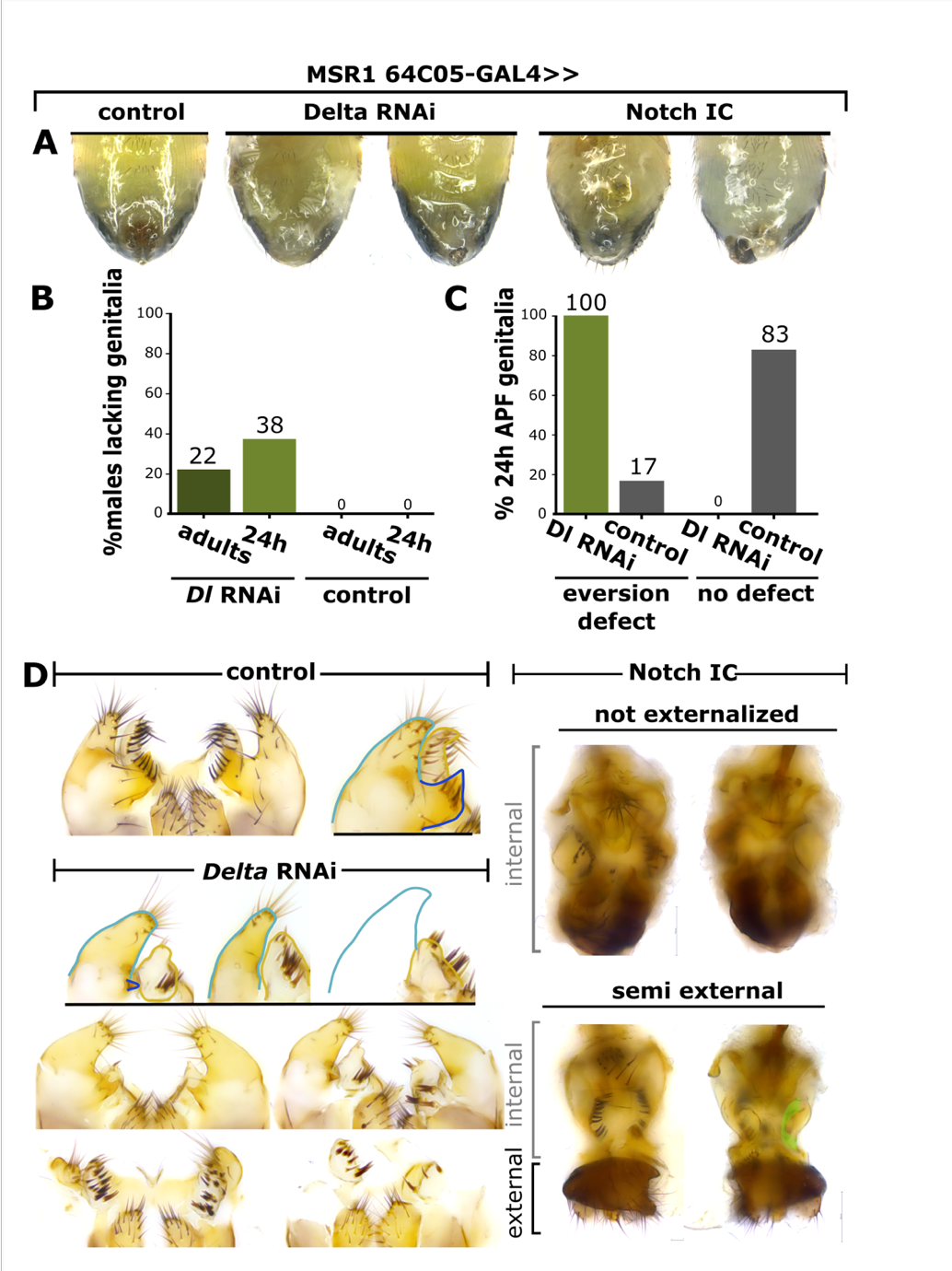

Fig. S11.

**phenotypes of early reduction of Delta by RNAi** (A-C) animals with severe genital eversion defects fail to develop genitalia. (A) Adult abdomens of control (left) Delta RNAi (middle), and Notch over expression(right). Abdomens from Delta RNAi and Notch overexpression display lack of external genitalia. (B) 22% of Delta RNAi adults and 38% of pupae dissected at 24h APF lack a recognizable genitalia. (C) 100% of Delta RNAi animals display a defect in genital eversion at 24h APF. (D) A range of adult genital defects represented in Delta RNAi genitalia that complete eversion. Over-activation of the Notch pathway in the L3 genital disc causes severe eversion defects. Represented adult genital cuticles were dissected from inside the abdomen, as they failed to externalize. An adult genitalia that has partially externalized from the abdomen resembles a wildtype developing pupal genitalia undergoing early stages of eversion (green shading).

Fig. S12.

Dumpy is deposited near Delta expressing cells at early stages. (A) Dumpy is deposited in an organized manner in the early everting pupal genitalia. A 21h APF reporter of Dumpy:YFP (green) stained with antibodies against Delta (grey) and Ecadherin (magenta). Top panel displays full projection of z-stacks, bottom panel displays a partial stack to visualize the inside of the everting pupal genitalia. Dumpy appears to be tethered to the Delta expressing epithelial folds (yellow arrowhead and white bracket, respectively). (B) Knockdown of Delta in the L3 genital disc also reduces Dumpy:YFP surrounding the male genital primordium bilaterally (bottom, compared to control). Reduction of Delta and Dumpy are indicated by white and yellow asterisks, respectively.

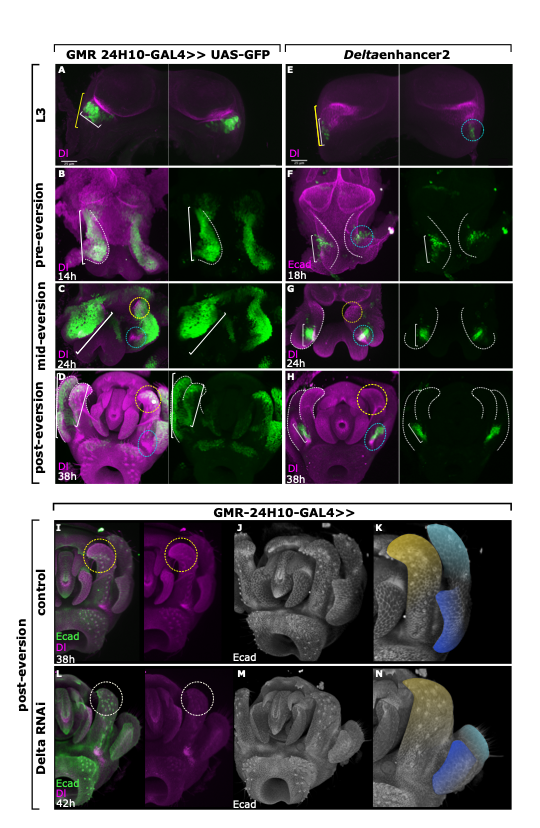

Fig. S13.

**The posterior lobe patterning signal emerges from a portion of a broader Delta pattern in the larval genital disc.** Activity of a larval enhancer of Delta (GMR24H10) (A-D) compared to the activity of Delta enhancer2 at four developmental stages (E-H). GFP marks reporter activity (white bracket), and Delta antibody is shown in magenta (yellow bracket in A and E). (A) GMR24H10 drives GFP activity in the lateral folds of the larval disc and overlaps Delta expression in this region. (B) At a pre-eversion stage, this activity is observed in a broad region that corresponds to the lateral plate/clasper (LP/CL) precursor (white bracket). (C) Mid-eversion, the LP/CL activity of the reporter overlaps the ventral expression of Delta at the tip of the future clasper (yellow dotted circle) but excludes the posterior lobe associated Delta pattern (blue dotted circle). (D) At a later stage post-eversion, GFP corresponds to the more ventral regions of the lateral plate and clasper, and does not overlap the lobe-associated pattern of Delta (blue dotted circle). (E) While Delta enhancer2 drives activity in the lateral folds of the larval genital disc (left:white bracket, right: blue dotted circle), this pattern overlaps only a portion of Delta’s endogenous expression domain (yellow bracket). (F) At a pupal stage pre-eversion, enhancer2 is only active in a subset of evaginating cells of the future LP/CL (white bracket and blue dotted circle). (G) Mid-eversion, enhancer2 only overlaps the lobe-accosiated expression of Delta (blue dotted circle) and does not overlap the ventral LP/CL activity of Delta (yellow dotted circle). (H) This is also observed post-eversion after the LP and CL have differentiated. (I-N) Knockdown of Delta using the GMR24H10-GAL4 driver affects the clasper and lateral plate (compare I-E with L-N). (I) In the control, Delta is expressed at the ventral tip of the clasper (yellow dotted circle), whereas knockdown of Delta strongly reduces this pattern (L, white dotted circle). (M, N) 3D rendering of post-eversion Delta-RNAi demonstrates defects in clasper and lateral plate development (compare to J and K). Claspers (false colored in yellow) of Delta-RNAi animals are shorter and less curved, and the lateral plates (false colored in light blue) are truncated.

Fig. S14.

**The posterior lobe associated localization of Dumpy is downstream of Delta.**

(A-C) Dumpy (green) is localized on the lateral plates in the lobe-forming region (yellow arrowheads in A and C) adjacent to Delta expressing cells (magenta, white bracket in A and B). (D-F) Knockdown of Delta reduces Dumpy:YFP deposition on the lateral plates (compare to control). Reduction of Delta and Dumpy are indicated by white and yellow asterisks, respectively. White boxes indicate resliced regions shown at the bottom of each panel. LP denotes the lateral plate, and C denotes the clasper.

Fig. S15.

**The spatial expansion but not the ancestral expression of Delta is downstream of Poxn.** Immunohistochemistry of Delta antibody in *D. melanogaster* in wildtype (A) and a Poxn null (Poxn^∆M220B5^) background (B-E). Flies mutant for Poxn do not exhibit an expanded expression pattern of Delta throughout pupal development. Arrows mark Delta expression at the base of the lateral plate/clasper associated with the posterior lobe. Poxn null allele described in (Boll & Noll, 2002)

| **transgenic lines** | **notes** | **notes, location** | **notes** | **source** |
| --- | --- | --- | --- | --- |
| enhancer2 Large∆ | 5548 bp Dl enhancer2 CRISPR | deletion over 3rd chr | balanced over TM6B | this study |
| Dl enhancer2-GAL4 #2 | Dl enhnacer2 GAL4 driver | attp40 |  | this study |
| Dl enhancer2-GAL4 #3 | Dl enhnacer2 GAL4 driver | attp40 |  | this study |
| BL45491 (Dl)GMR-23G03-GAL4 | Dl L3 enhanecer | immediately upstream | L3 overlap w/Dl, weak signal | Bloomington Drosophila stock center |
| BL49063 (Dl)GMR-24B08-GAL4 | Dl L3 enhanecer | Dl intron1 | L3 overlap w/Dl, specific | Bloomington Drosophila stock center |
| BL45505 (Dl)GMR-24H10-GAL4 | Dl L3 enhanecer | 1351bp US, overlaps 23G03 | L3 overlap w/Dl, strong | Bloomington Drosophila stock center |
| BL46170 (Dl)GMR-25B11-GAL4 | Dl L3 enhanecer | Dl intron1 | L3 overlap w/Dl, strong & broad | Bloomington Drosophila stock center |
| BL49104 (Dl)GMR-25A03-GAL4 | Dl L3 enhanecer | Dl intron2 | L3 overlap w/Dl, weak signal | Bloomington Drosophila stock center |
| BL49375 (Dl)GMR-37A05-GAL4 | Dl L3 enhanecer | Dl intron3 | L3 overlap w/Dl, strong & broad | Bloomington Drosophila stock center |
| BL49600 (MSR1)GMR64C05-GAL4 | target Dl L3 activity in the lateral folds |  |  | Bloomington Drosophila stock center |
| w;dpyYFP/cyo; MSR1-64C05-G4/TM6B | dpy:YFP crossed into L3 driver | visualize dpy in a Dl RNAi bg | select red eyes | in house |
| UAS-NotchIC/cyo.GFP (FBtp0012228) | Notch Intra-cellular domain | constitutively active Notch |  | personal gift from Gerard Campbell |
| UAS-Delta RNAi (28032) |  |  |  | Bloomington Drosophila stock center |
| UAS-mCherry RNAi (35785) | control |  |  | Bloomington Drosophila stock center |
| UAS-GFP (1522) |  |  |  | Bloomington Drosophila stock center |
| Dumpy:YFP (115238) |  |  |  | Drosophila Genomics and Genetic Resources |

Table S1.

**Fly stocks used in this study**

| **Description** | **Sequence** | **Notes** |
| --- | --- | --- |
| *D. melanogaster* *Delta* enhancer1 forward primer | TTCCGggcgcgccTGCGAATTGCTTGTACGTGACACG | Lrg F (LrgA) |
| *D. melanogaster* *Delta* enhancer1 reverse primer | TTGCCcctgcaggATCGACACGCTTAACTGGCA | Lrg R4 (LrgA) |
| *D. ananassae* *Delta* enhancer1 forward primer | TTCCGggcgcgccTGCGAATTGCTTGTACGTGACACG |  |
| *D. ananassae* *Delta* enhancer1 reverse primer | TTGCCcctgcaggATCGACACGCTTAACTGGCA |  |
| *D. biarmipes* *Delta* enhancer1 forward primer | TTCCGggcgcgccTGCGAATTGCTTGTACGTGACACG |  |
| *D. biarmipes* *Delta* enhancer1 reverse primer | TTGCCcctgcaggCRTTGCGTGTAATTAAAACRCMARC |  |
| *D. melanogaster* *Delta* enhancer2 forward primer | TTCCGggcgcgccACCAGGGATGCAGCCTAGTTC |  |
| *D. melanogaster* *Delta* enhancer2 reverse primer | TTGCCcctgcaggTTCGCTGTCCGAGAATGTCAAC | FR3 |
| *D. ananassae* *Delta* enhancer2 forward primer1 | TTCCGggcgcgccTAGCCCAACAACAGGCGGAAGAT |  |
| *D. ananassae* *Delta* enhancer2 reverse primer1 | TTGCCcctgcaggCCCTCTGGCATATATGTATAGC |  |
| *D. biarmipes* *Delta* enhancer2 forward primer1 | TTCCGggcgcgccCACCCACAGAGGTACCACGTATA |  |
| *D. biarmipes* *Delta* enhancer2 reverse primer1 | TTGCCcctgcaggTACTCCGCTGTCCAAGTGTGTC |  |
| *D. melanogaster Delta* probe forward primer | ACTCTTGGTGCAGCAGGTACTG |  |
| *D. melanogaster* *Delta* probe reverse primer | taatacgactcactataggGAGTGTCCAAATGAATCGTCGC |  |
| *D. ananassae Delta* probe forward primer | TCGCTGGAGTACGATTTCCG |  |
| *D. ananassae Delta* probe reverse primer | taatacgactcactataggTGCCATCCGGTCAAACAGAT |  |
| *D. biarmipes Delta* probe forward primer | TCGCTGGAGTACGATTTCCG |  |
| *D. biarmipes Delta* probe reverse primer | taatacgactcactataggTGCCATCCGGTCAAACAGAT |  |
| *mbeta* reporter forward primer | TTCCGggcgcgccTGGGAACTGGACAGTGAGTG |  |
| *mbeta* reporter reverse primer | TTGCCcctgcaggGGTCCAGCGGACAAAACGTT |  |
| *mbeta* probe forward primer | GACCTATCAGTACCGCAAG |  |
| *mbeta* probe reverse primer | taatacgactcactatagggagaCGCAGCTTCTTCATGTGCTC |  |
| Primer for synthesizing sgRNA targeting the left homology arm of enhancer1 (sense) | **GTCG**CCCTTAATCTGGACACTAAT |  |
| Primer for synthesizing sgRNA targeting the left homology arm of enhancer1 (antisense) | **AAAC**ATTAGTGTCCAGATTAAGGG | gRNA left B |
| Primer for synthesizing sgRNA targeting the right homology arm of enhancer1 (sense) | **GTCG**AGACTTAGCCGGTGGGGATG |  |
| Primer for synthesizing sgRNA targeting the right homology arm of enhancer1 (antisense) | **AAAC**CATCCCCACCGGCTAAGTCT | gRNA right B |
| Forward primer for cloning the left homology arm to target enhancer1 | TTCCGggcgcgccGAACCTCACATATGAACCCATC | Left AB F |
| Reverse primer for cloning the left homology arm to target enhancer1 | TTGCCcctgcaggAGTGTCCAGATTAAGGGATATC |  |
| Forward primer for cloning the right homology arm to target enhancer1 | TTGCCccgcggCCCCACCGGCTAAGTCTTTCAC |  |
| Reverse primer for cloning the right homology arm to target enhancer1 | TTCCAagatctTGCTTTGTCTCGTCTCGTCTCG |  |
| Primer for synthesizing sgRNA targeting the left homology arm of enhancer2 | aaaataatacgactcactataggGACTTGAATGTTCCGTTGTTgttttagagctag | T9Large_L1_gRNA1 |
| Primer for synthesizing sgRNA targeting the right homology arm of enhancer2 | aaaataatacgactcactataggAGGCAGTCAGGTTGTTCACGgttttagagctag | T9Large_R1_gRNA |
| Forward primer for cloning the left homology arm to target enhancer2 | tcttgcatgctagcggccgcAACACCACCACCTGAACCAGCT | T9Large_HArmL_F |
| Reverse primer for cloning the left homology arm to target enhancer2 | gtgcatatgtccgcggccgcAACGGAACATTCAAGTCAGACCC | T9Large_HArmL1_R |
| Forward primer for cloning the right homology arm to target enhancer2 | ctccatgcataaggcgcgccACGGGGTTAAACGGTTGTTGAC | T9Large_HArmR1_F |
| Reverse primer for cloning the left homology arm to target enhancer2 | gcagaaggcctaggcgcgccCTATGAGTTCCACTGGGCTGAC | T9Large_HArmR_R1 |
| pHD-dsRed_rightF_dns1 | GACTACACCATCGTGGAGCAGT | enhancer2 CRISPR validation primers |
| pHD-dsRed_leftR_dns1 | TGAACTCCTTGATGACGTCCTC | enhancer2 CRISPR validation primers |
| Tile9_Right_outArm_R | ATTGACGCTGACAAGGCTTGGC | enhancer2 CRISPR validation primers |
| Tile9_Right_inArm_R | GCAATTTCGCCCTTGGCAATTG | enhancer2 CRISPR validation primers |
| Tile9_Left_inArm_F | GGCTTAGATGCACAGTGGTA | enhancer2 CRISPR validation primers |

Table S2.

**Primer oligonucleotides used in this study**
